# The HMGB1-RAGE axis modulates the growth of autophagy-deficient hepatic tumors

**DOI:** 10.1101/847673

**Authors:** Bilon Khambu, Honghai Hong, Sheng Liu, Gang Liu, Xiaoyun Chen, Zheng Dong, Jun Wan, Xiao-Ming Yin

**Author notes:** Correspondence to: Bilon Khambu, PhD Department of Pathology and Laboratory Medicine Indiana University School of Medicine, Indianapolis, IN 46202.

## Abstract

Autophagy is an intracellular lysosomal degradative pathway important for tumor surveillance. Autophagy deficiency can lead to tumorigenesis. Autophagy is also known to be important for the aggressive growth of tumors, yet the mechanism that sustains the growth of autophagy-deficient tumors is not known. We previously reported that progression of hepatic tumors developed in autophagy-deficient livers required high mobility group box 1 (HMGB1) that is released from autophagy-deficient hepatocytes. However, the mechanism by which HMGB1 promotes hepatic tumorigenesis is not understood. In this study we examined the pathological features of the hepatic tumors and the mechanism of HMGB1-mediated tumorigenesis using liver-specific autophagy-deficient (*Atg7-/-*) and *Atg7-/-/Hmgb1-/-* mice. We found that in *Atg7-/-* mice the tumors cells were still deficient in autophagy and could also release HMGB1. Histological analysis using cell-specific markers suggested that fibroblast and ductular cells were present only outside the tumor whereas macrophages were present both inside and outside the tumor. Genetic deletion of HMGB1 or one of its receptors, receptor for advanced glycated end product (*Rage*), retarded liver tumor development. In addition, we found that expression of RAGE was only on ductual cells and Kupffer’s cells but not on hepatoctyes, which suggested that HMGB1 might promote hepatic tumor growth through a paracrine mode that altered the tumor microenvironment. Furthermore, HMGB1 and RAGE enhanced the proliferation capability of the autophagy-deficient hepatocytes and tumors. Finally, RNAseq analysis of the tumors indicated that HMGB1 induced a much broad changes in tumors. In particular, genes related to mitochondrial structures or functions were enriched among those differentially expressed in tumors in the presence or absence of HMGB1, revealing a potential key role of mitochondria in sustaining the growth of autophagy-deficient liver tumors via HMGB1 stimulation.

## Introduction

Autophagy is an important mechanism regulating tumorigenesis. Its dysfunction due to external stress or genetic inactivation may lead to tumorigenesis. Indeed, liver-specific deletion of *Atg5* or *Atg7* causes spontaneous development of liver tumors ^1–4^. Similarly, reduced autophagy from constant activation of mammalian target of rapamycin complex 1 (mTORC1) also promotes hepatic neoplastic transformation. For example, deletion of phosphatase and tensin homolog (PTEN) or tuberous sclerosis complex 1 (TSC1) leads to constitutive activation of mTORC1, and a decrease in autophagic activity, causing spontaneous HCC ^5, 6^. These studies suggest that hepatocytes require the tumor-suppressive function of autophagy for maintaining its homeostasis.

Excessive reactive oxygen species (ROS) generated due to autophagy-deficiency is implicated in tumor development ^7, 8^. Consequently, pharmacological inhibition of ROS formation by the antioxidant N-acetylcysteine results in a strong suppression of tumor development in *Atg5*-deficient liver^8^. Moreover, there is a persistent activation of an anti-oxidative stress-related transcription factor NFE2L2/ NRF2(nuclear factor, erythroid 2 like 2) to limit the oxidative injury ^9^. Paradoxically, codeletion of *Nrf2* gene also prevents tumorigenesis in the autophagy-deficient liver ^1, 3^.

On the other hand, autophagy could also regulates hepatic tumorigenesis by modulating the release of a damage-associated molecular pattern (DAMP) molecule, high mobility group box 1 (HMGB1). Our previous work discovered that defective autophagy leads to the release of HMGB1, which promotes hepatic tumorigenesis ^2^. HMGB1 is an extensively studied non-histone nuclear protein that facilitates binding of regulatory proteins to DNA and typically enhances transcriptional activation ^10^. It is known that nuclear HMGB1 can be released into the extracellular environment and acts as an immune mediator in sterile inflammation. However, codeletion of *Hmgb1* in the autophagy-deficient liver results in delayed tumor development via a mechanism independent of its usual role in injury, inflammation, and fibrosis ^2^.It is thus unclear how HMGB1 mechanistically promotes hepatic tumorigenesis in autophagy-deficient liver.

In the present study, we have characterized the cellular and molecular context of the hepatic tumors driven by autophagy deficiency. We observed that the tumors were originated mainly from the autophagy-deficient hepatocytes that had already released HMGB1. Furthermore, we showed that HMGB1 and its dominant receptor RAGE positively affect the proliferation of tumor cells. RNA sequencing analysis identified expressional differences in multiple genes and signaling pathways in tumors derived from *Atg7-/-* liver and in tumors derived from *Atg7-/-/Hmgb1-/-* livers. Our data, therefore, identify a key role of HMGB1 in autophagy-deficient tumor growth. HMGB1 could thus be a potential therapeutic target.

## Results

### 1. Hepatic tumor cells in autophagy-deficient livers have features consistent with autophagy deficiency

Autophagy possesses both antitumorigenic and protumorigenic role, depending on whether it occurs before or after the onset of tumorigenesis. Autophagy-deficient livers develop tumors, confirming the surveillance role of autophagy in the liver. The tumor first appears at the 9-month of the age and the tumor size and the number gradually increase as the mice get older^2, 3^. The tumors in *Atg5*- or *Atg7*- conditional knock out livers seem to be hepatic adenoma, which does not progress to carcinoma or metastasis stage^3^. However, the molecular and cellular nature of these tumors had not been fully characterized.

We thus examined hepatic tumor from autophagy-deficient mice(*Atg7-/-*) mice. Lack of the ATG7 expression was confirmed in tumor and non-tumor liver tissue from *Atg7-/-* mice when compared to age-matched *Atg7 F/F* mice (**Figure 1A**). The level of the autophagy substrate SQSTM1 was also higher in *Atg7-/-* tumor as in the non-tumor tissue. Analysis of LC3B, an autophagy-specific marker showed no formation of LC3B-II in the *Atg7-/-* tumor as in the non-tumor samples (**Figure 1A**). These results indicated that the tumor tissues were also deficient in autophagy and thus they would have arisen from the autophagy-deficient hepatocytes. We further confirmed this notion by examining the expression of SQSTM1 and ubiquitin(UB) in the liver. Immunohistological and immunofluorescence analysis was performed by taking images of eight different regions covering the non-tumor, peri-tumor, and the tumor regions as shown in **Figure 1B**. A clear accumulation of SQSTM1 and UB in the tumor region of the autophagy-deficient liver was observed, which was at the level similar to that in the non-tumor tissues (**Figure 1C-D**), suggesting that the tumor tissues were defective in autophagy and had defective protein quality control. In addition, the tumor tissues were all positive for the hepatocyte-specific marker, HNF4α, which were colocalized in the same cells that had elevated SQSTM1 and UB staining (**Figure 1F**).

**Figure 1:**
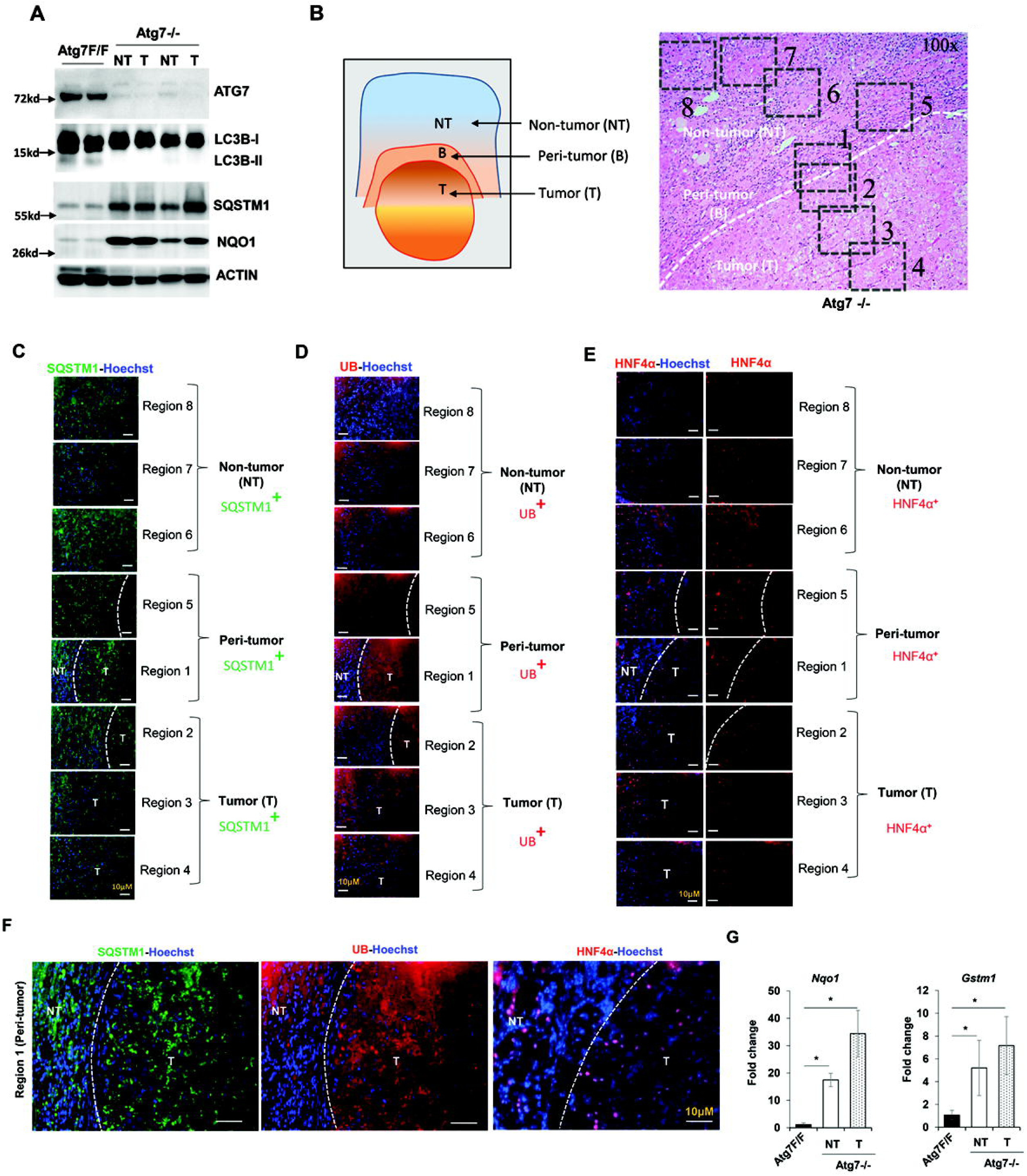
Hepatic tumor in autophagy-deficient livers are derived from autophagy-deficient hepatocytes. (**A**) Immunoblot analysis of autophagy function-related proteins (ATG7, SQSTM1, LC3B-I/II) and NRF2 pathway-related proteins(NQO1) in whole livers isolated from 15-month old *Atg7F/F*, and *Atg7-/-* mice. (**B**) Schematic representation of the non-tumor, peri-tumor, and tumor region of the liver sections. Region 1 and Region 5: peri-tumor region, Region 2-Region 4: tumor region, and Region 6-Region 8: non-tumor region. (**C-E**) Livers from 12-month old mice of *Atg7-/-* genotype were sectioned and immunostained with anti-SQSTM1(C), Anti-Ubiquitin (UB) (D), or anti-HNF4_α_ (E). Dotted lines indicate the tumor border. (**F**) Magnified image of the region 1(peri- and intra-tumor region) of panels C, D, & E. (**G**) The hepatic mRNA expression level of NRF2 target genes, *Nqo1 and Gstm1,* in the livers of 15-month old *Atg7F/F*, and in the non-tumor and tumor samples from the liver of age-matched *Atg7-/-* mice. NT, non-tumor, T, tumor. Data are reported as mean± SE,* *P*<0.05; n=3 mice per group.

We next analyzed whether the accumulation of SQSTM1 in tumor tissue could activate the anti-oxidative response-related NRF2 transcription factor as in non-tumor tissues ^1, 9^. We found that one NRF2 target protein, NQO1, were drastically elevated in the tumor tissues of the *Atg7-/-* mice (**Figure 1A**). The mRNA level of the NRF2 target genes, *Nqo1* and *Gstm1* were also significantly elevated in the *Atg7-/-* samples whether they were from non-tumor or tumor tissue (**Figure 1G**). These observations indicated that hepatic tumors in autophagy-deficient livers arise from the autophagy-deficient hepatocytes with alterned NRF2 and SQSTM1 levels.

### 2. Hepatic progenitor cells are localized exclusively in the non-tumor region but not inside the tumor

Hepatic progenitor cells(HPC), also known as oval cells or ductular cells, expand during chronic liver injury in patients and in rodents^11, 12^. The expansion of HPCs is significant in the autophagy-deficient livers ^2^. HPC has been noted to possess the capacity to become tumorigenic in vivo when transduced with H-ras and SV40LT^13^. We thus explored the relationship of these cells to the tumor in autophagy-deficient livers by examining their spatial interactions.

H-E staining showed that the distribution of HPCs was mostly around the tumor-adjacent region (**Figure 2A**). In the area of tumor tissues, the normal tissue architecture, such as bile duct, and portal tract formation, was completely lost. Moreover, the tumor region was composed of irregular hepatic plates with tumor cells showing large nuclear-cytoplasmic ratio and occasionally nuclear atypia (**Figure 2A**). Immunostaining for CK19, a common marker for expanded HPC showed that the hepatic tumors were negative for CK19 (**Figure 2B**). Instead, most of the CK19 positive cells appear to form a compact sheet surrounding the tumor (**Figure 2B**). Further analysis of the HPC distribution using Sox9 as another marker revealed a similar pattern of distribution exclusively in the peritumor and non-tumor regions (**Figure 2C**). Some of the HPCs were positive for SQSTM1 aggregates although many did not show elevated SQSTM1(**Supplementary Figure S1A-B**). The possibility that some of this SQSTM1 positive HPC may be derived from the autophagy-deficient hepatocytes cannot be excluded as such transdifferentiation had been reported previously ^11, 14^.

**Figure 2.**
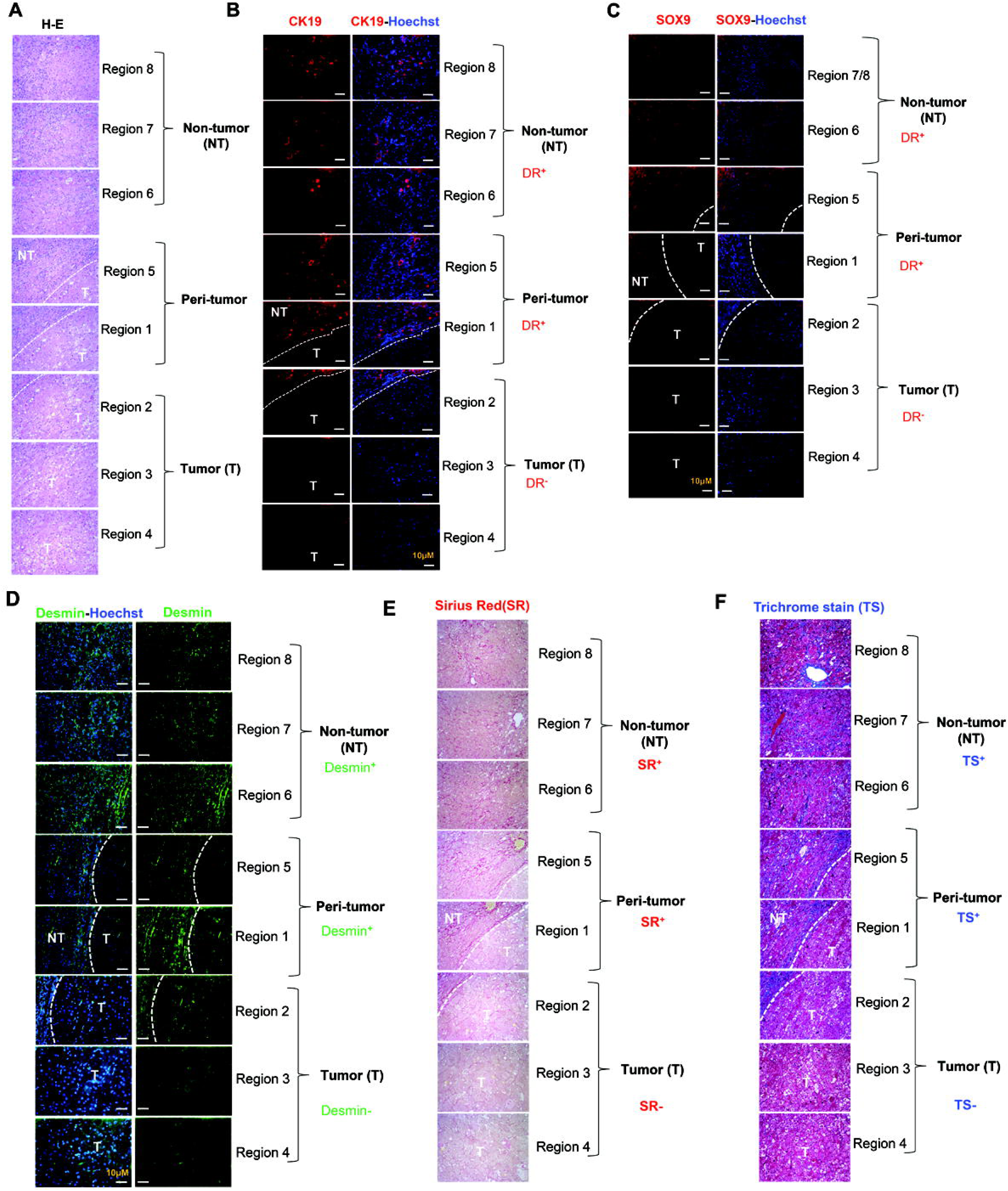
Hepatic Progenitor Cells and fibrosis are localized exclusively in peri-tumor and non-tumor regions but are absent inside the tumor. Liver sections from 12-month old mice of the *Atg7-/-* genotype were subjected to H-E staining (**A**) (original magnification, X200) and immunostaining for CK19 (**B**), SOX9 (**C**), Desmin immunostaining (**D**), Sirius Red stain (**E**), or Trichrome stain (**F**) (original magnification, X200). Dotted lines indicate the tumor border. NT, non-tumor, T, tumor.

Interestingly, HPC and liver cancer stem cells (CSC) also share several cellular markers^15^. Markers such as EpCAM, CD133, and CD24 have been used for isolating CSC with stem cell features ^15, 16^. HPC in the context of chronic liver injury has also been considered as one possible origin of liver CSC. We thus analyzed the expression of these CSC markers in the non-tumor and tumor tissues of the autophagy-deficient liver. Real-time PCR analysis showed that the expression of *Cd133*, *Cd200*, *Cd34*, *Cd44*, *Ly6a/Sca1*, and *Ly6d* were significantly upregulated in *Atg7-/-* liver tissues compared to control *Atg7 F/F* mice (**Supplementary Figure S2A**). The elevation of these CSC markers in the tumor tissues also suggested that tumors have a precursor/stem-cell phenotype. Such induction was not observed with *Cd24a*, and *Cd90,* suggesting the possible heterogeneity in the CSC in the tumor and non-tumor tissues of *Atg7-/-* mice (**Supplementary Figure S2A**). Interestingly, most of the stemness-related transcription factors such as *Oct4*, *Nanog*, *Klf4* and *Sox2* were significantly downregulated in *Atg7-/-* livers as compared to *Atg7F/F* livers (**Supplementary Figure S2B**). The lack of expression of Nanog has been linked to the adenoma nature of the tumor ^17^. These changes were not more significant in the tumor tissue than in the non-tumor tissues, and thus may not be the mechanisms discriminating the two types of tissues.

HPCs that are mostly detected in peritumoral areas has been reported to express multiple angiogenic paracrine factors such as vascular endothelial growth factor(VEGF), platelets-derived growth factor(PDGF), and angiopoietin (ANGPT) in pediatric hepatoblastoma and HCC ^18^. These HPCs could interact with pro-tumorigenic cells heterotypically via mitogenic factors. We thus examined the expression of several angiogenic markers. Real-time PCR analysis indicated that expression of angiogenic factor *Angpt2* and *Pdgfb* were significantly upregulated in liver tumor and non-tumor tissues compared to wild-type *Atg7 F/F* mice (**Supplementary Figure S3**). Such induction was not observed with *Vegfa* and *Angpt1* (**Supplementary Figure S3**), suggesting the possible heterogeneity in the angiogenic factors acting within the peritumoral niche of *Atg7-/-* mice. Taken together, the distinct separation of the HPC and tumor cells in the *Atg7-/-* liver suggests that the HPC may not evolve into the tumor cells but could contribute to a tumor microenvironment that affects the tumor development.

### 3. Fibrosis is present in the peri-tumor region and encapsulates the tumor

Development of hepatic tumors are strongly associated with the status of liver fibrosis, with 80-90% of HCCs developing in the fibrotic or cirrhotic livers ^19^. On a cellular level, fibrogenesis is most significantly mediated by the activation of hepatic stellate cells (HSCs) that transdifferentiate from Vitamin A-storing pericyte-like cells to alpha-smooth muscle actin (α-SMA)-positive, collagen-producing myofibroblasts in response to liver injury. Since liver fibrosis is one of the earliest events that occur in the autophagy-deficient liver ^2^, we examined how closely the tumor cells were associated with liver fibrosis.

Immunostaining analysis indicated that the number of desmin-expressing HSCs was remarkably increased in *Atg7-/-* mice (**Figure 2D**). Unlike the distribution of macrophages, the desmin positive HSC were absent inside the tumor but were present in the non-tumor and peri-tumor regions of the liver (**Figure 2D**). Consistent with the increased desmin positive HSCs in peri-tumor and non-tumor region, increased fibrillar collagen deposition was detected by Sirius Red and Trichome stain in the non-tumor and peri-tumor region (**Figure 2E-F**). Collagen deposition was notably absent inside the tumor of the autophagy-deficient liver (**Figure 2E-F**). Taken together, the peri-tumoral desmin positive HSCs may be responsible for the production of the fibers that encapsulated and demarcated the tumor tissue. It is possible that fibrosis in the autophagy-deficient liver may play an inhibitory role against tumor infiltration into normal tissues, thus contributing to the more benign presentation of the tumorigenesis in this setting.

### 4. Macrophages but not other immune cells can be found inside the tumor tissue

Hepatocellular neoplasia often occurs in the setting of chronic injury and inflammation, which is present in autophagy-deficient livers ^2, 20^. Persistent inflammation is also known to promote and exacerbate malignancy. Among many different types of inflammatory cells, the tumor-associated macrophage (TAM) is a key component involved in the initiation and maintenance of tumor cells ^21^. Factors secreted by the TAM are thought to contribute to the initiation and promotion of tumors. Since the identification of inflammatory and immune cells assists in characterizing the nature of the hepatic tumor and their potential contribution to the tumor growth, we next examined the distribution of inflammatory and immune cells in the tumor-bearing *Atg7-/-* mice.

Immunohistological staining for the hepatic kupffer’s cells (F4/80^+^) showed their presence in both tumoral and non-tumoral regions (**Figure 3A**). In contrast, most of the myeloperoxidase (MPO)-positive neutrophils, CD3-positive T cells, and CD45R-positive B cells were absent from the tumoral region but present exclusively in the non-tumor region (**Figure 3B-D**). Quantitative RT-PCR also found a strong upregulation of F4/80 and Ly6c expression in 12-month old *Atg7-/-* livers as compared to age-matched *Atg7F/F* livers, and there was a further elevation in tumor tissues (**Figure 3E**). The CD4 mRNA level was modestly elevated but the CD8 mRNA level was significantly suppressed in the tumor-bearing *Atg7-/-* liver (**Figure 3E**).

**Figure 3.**
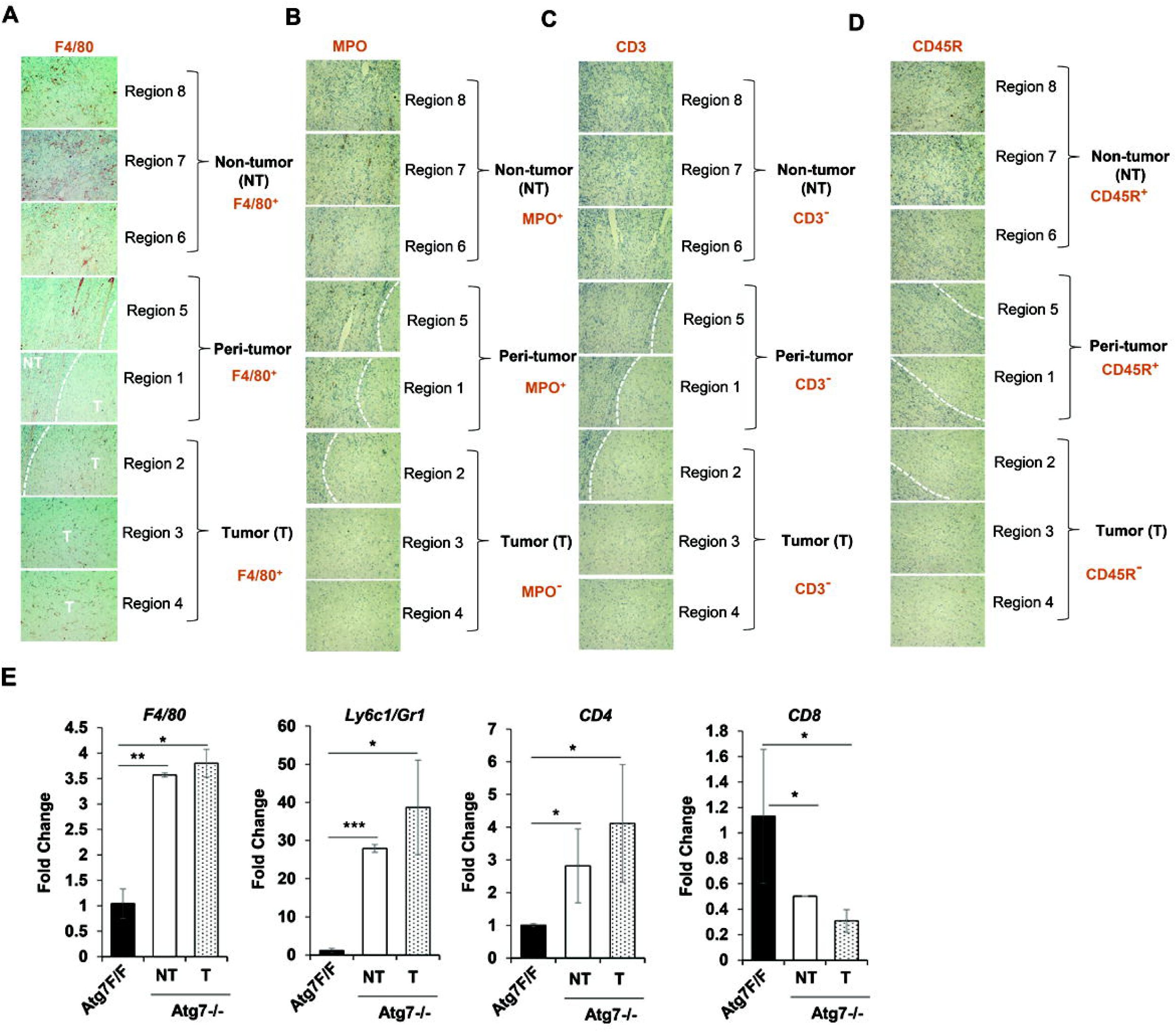
Macrophages but not other immune cells are found within the tumor. Liver sections from 12-month old mice of *Atg7-/-* genotype were subjected to immunohistochemistry staining for F4/80 (**A**), Myeloperoxidase (MPO) (**B**), CD3 (**C**) and, CD45R (**D**) (original magnification, X100). Dotted lines indicate the tumor border. (**E**) The hepatic mRNA expression level of immune cell-associated genes in 15-month old *Atg7F/F* and *Atg7-/-* liver tissues. NT, non-tumor, T, tumor. Data are reported as mean± SE,* *P*<0.05, ** *P*<0.01, *** *P*<0.001, n.s.: no significance; n=3 mice per group.

Macrophages can play important roles in regulating hepatocytes proliferation and survival by secreting cytokines such as tumor necrosis factor-α(TNFα), IL-6, and IL-1β and growth factors such as VEGF, hepatocytes growth factors(HGF) and transforming growth factors(TGFs). TNFα produced by infiltrating macrophages could activate TNF receptor 1(TNFR1)-NF-kB signaling in hepatocytes, resulting in enhanced tumor growth ^22^. In contrast to the presence of infiltrating F4/80 macrophages and elevated expression of F4/80 and Ly6c, the mRNA expression of a set of inflammatory cytokines such as *TNF*α, *IL-6*, *Il-1*β, and *IL-17* were strongly downregulated in the 12-month old tumor-bearing *Atg7-/-* liver (**Supplementary Figure S4**). These data suggest that there is ongoing non-resolving inflammation in tumor and non-tumor tissue of autophagy-deficient mice but their contribution to tumor growth has yet to be fully determined.

### 5. Autophagy deficient hepatic tumor cells release HMGB1

Autophagy-deficient livers manifest multiple pathological changes, including liver injury, inflammation, fibrosis, ductular reaction and tumor development ^1, 2, 23^. Moreover, the autophagy-deficient hepatocytes continuously release HMGB1, an intracellular nuclear DAMP protein, to impact the expansion of HPC ^2^. Given that HMGB1 as a secretory factor could recruit other hepatic cells such as inflammatory cells or fibrotic cells to the tumor, favoring the buildup of permissive microenvironment ^24, 25^, we sought to determine whether the tumor tissues also release HMGB1 similar to the non-tumor autophagy-deficient liver tissue.

As anticipated, the immunoblot analyses found that less HMGB1 proteins are present in tumor and non-tumor tissue of the *Atg7-/-* liver, as compared to the *Atg7 F/F* liver (**Figure 4A**). Co-immunofluorescence staining of HMGB1 and SQSTM1 also showed that the tumor cells with the accumulated SQSTM1 were devoid of both nuclear and cytosolic HMGB1 (**Figure 4B**). The mRNA level of HMGB1 was comparable between the liver tissues of *Atg7F/F*, and *Atg7-/-* mice (**Figure 4C**). Thus, the loss of HMGB1 protein in hepatic tumor cells supports further the notion that the nature of these cells are autophagy-deficient, and suggests that these tumor cells had released HMGB1, which could affect the microenvironment.

**Figure 4.**
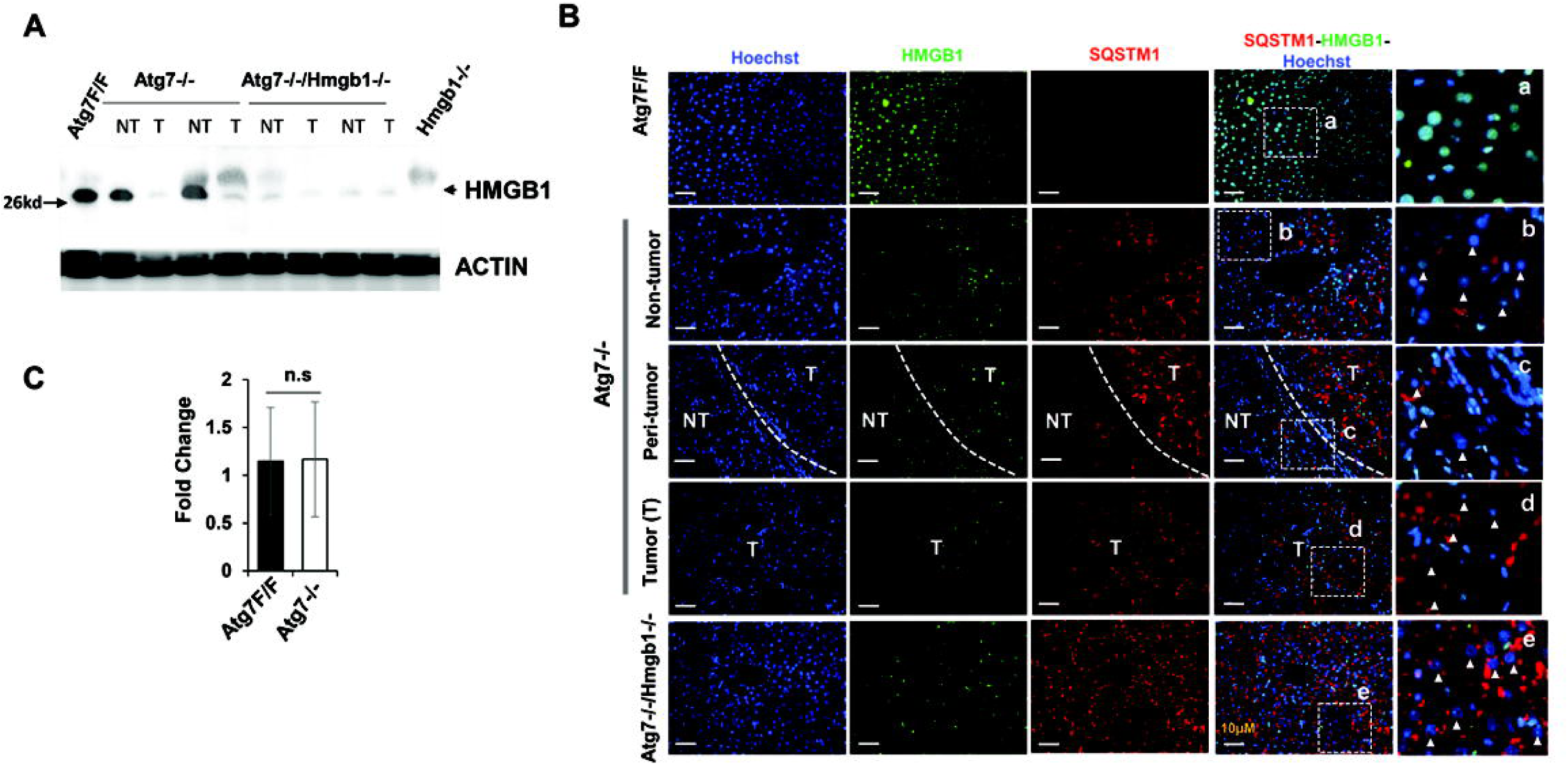
Hepatic HMGB1 is absent in the tumor of autophagy-deficient livers. (**A**) Livers of 15-month old mice of different genotypes were examined for HMGB1 by immunoblotting assay. (**B**) Liver sections from 15-month old mice of different genotypes were immunostained with anti-HMGB1 and anti-SQSTM1. White dotted lines indicate the tumor border. White arrowhead indicates the hepatocytes without nuclear HMGB1. (**C**) The hepatic mRNA expression level of *Hmgb1* in 15-month old *Atg7F/F* and *Atg7-/-* mice, determined by real-time PCR. NT, non-tumor, T, tumor. Data are reported as mean± SE, n.s., no significance; n=3 mice per group.

### 6. HMGB1 promotes hepatic proliferation

HMGB1 has a mitogenic effect in many cell lines including in vitro human HCC cell lines ^26^. HMGB1 released by autophagy-deficient hepatocytes could affect the growth of tumorigenic hepatocytes, thus promoting hepatocarcinogenesis. Indeed we had found that HMGB1 was important for the tumorigenesis in the autophagy-deficient liver ^2^.

Consistently, we now found that *Atg7-/-* livers had a remarkably increased number of the hepatocytes positive for Proliferation of cell nuclear antigen (PCNA) (**Figure 5A**). PCNA positive cells seem to be present in both non-tumor and tumor regions without much differences in the level. This observation was confirmed by immunostaining for Ki67 positive cells (**Supplementary Figure S5A-B**). Ki67 positive cells were elevated in non-tumor and intra-tumor regions of the *Atg7-/-* liver without differences in the number of proliferating cells (**Supplementary Figure S5A**).

**Figure 5.**
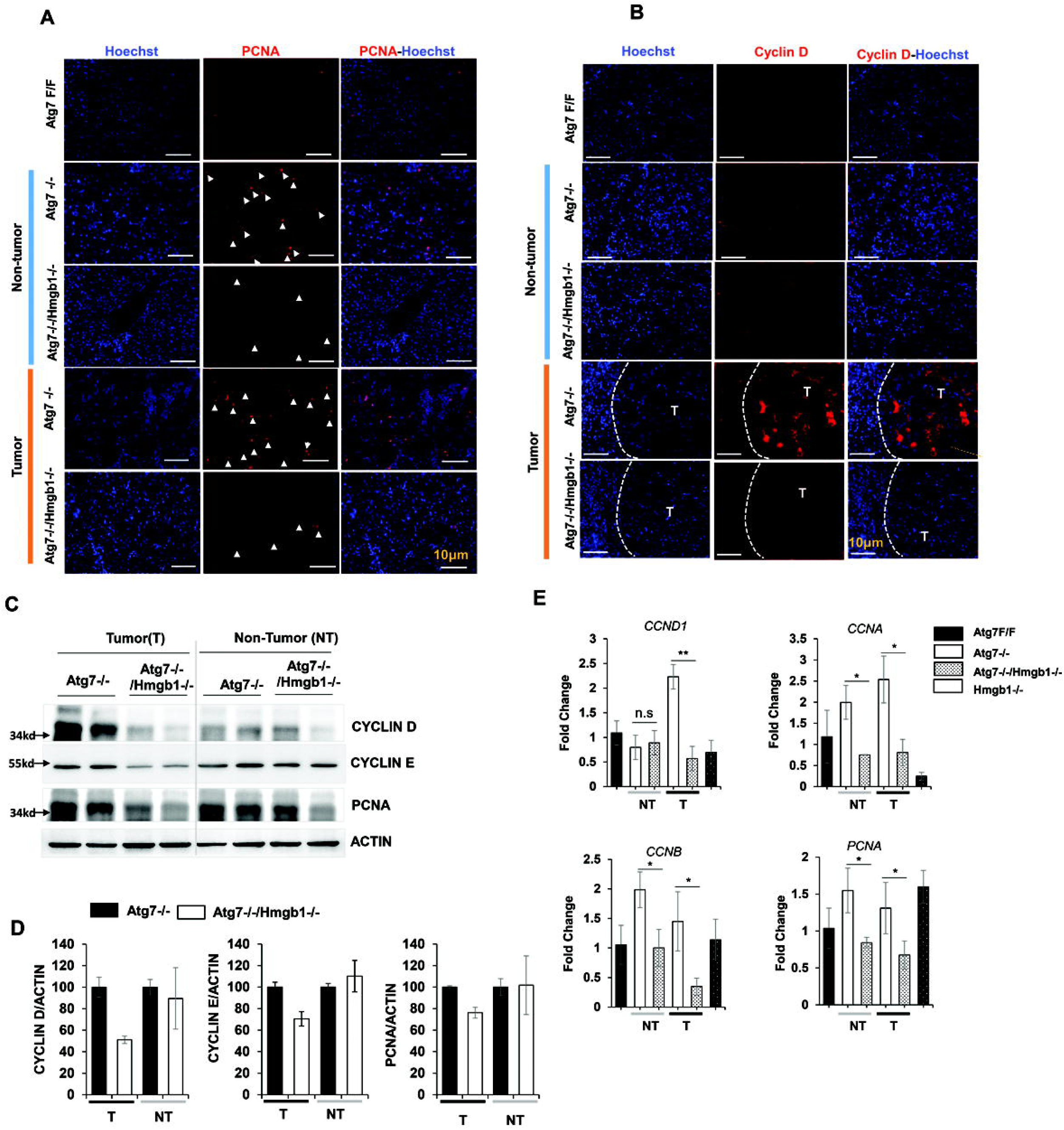
Loss of HMGB1 in hepatocytes correlates with reduced proliferation in the tumor. (**A-B**) Liver sections from 15-month old mice of different genotypes were immunostained with anti-PCNA(A), or anti-Cyclin D (B). White arrow indicated proliferating hepatocytes. White dotted lines indicate the tumor border. (**C**) Immunoblot analysis of PCNA, cyclin D1, and cyclin E proteins in the tumor or non-tumor sample of 15-month old *Atg7-/-* and, *Atg7-/-/Hmgb1-/-* mice. (**D**) Densitometry qualification of the indicated proteins. (**E**) The hepatic mRNA level of indicated genes were determined in the indicated tissues of 15-month old mice of different genotypes, determined by real-time PCR. NT, non-tumor, T, tumor. Data are reported as mean± SE,* *P*<0.05, ** *P*<0.01, n.s., no significance; n=3 mice per group.

To determine the role of HMGB1 in the proliferation of hepatocyte, we compared the cellular proliferation in 15-month old *Atg7-/-* and *Atg7-/-/Hmgb1-/-* mice because both genotypes developed a notable but different number of tumors at this age ^2^. Immunostaining analysis for PCNA showed a lower number of proliferating hepatocytes in the tumor of *Atg7-/-/Hmgb1-/-* livers than those in the *Atg7-/-* livers (**Figure 5A**). Interestingly, the number of PCNA positive cells was also lower in the non-tumor region of *Atg7-/-/Hmgb1-/-* livers when compared to the non-tumor region of *Atg7-/-* livers (**Figure 5A**).

Because the increased cellular proliferation is associated with increased expression of cyclin protein, we next examined the expression of Cyclins. First, immunostaining found that the expression of Cyclin D1 was more up-regulated in the tumor samples of *Atg7-/-* liver than in the tumors of the *Atg7-/-/Hmgb1-/-* livers (**Figure 5B**, **Supplementary Figure S6**). Second, immunoblot analysis of Cyclin E, similar to PCNA protein, also showed a higher level in tumor and non-tumor regions of the *Atg7-/-* livers than that in the *Atg7-/-/Hmgb1-/-* livers (**Figure 5C-D**). Real-time PCR analysis demonstrated that hepatic expression of *CCND1*, *CCNA1*, and *CCNB1* were significantly up-regulated in *Atg7-/-* mice, compared to *Atg7 F/F* mice(**Figure 5E**). The expression of *CCND1* and *CCNA1* was even more pronouncedly elevated in the tumor region than in the non-tumor tissues from *Atg7-/-* mice (**Figure 5E**). Such induction was not observed in tumor tissues from *Atg7-/-/Hmgb1-/-* mice (**Figure 5E**), suggesting that *Hmgb1* deletion retarded cell cycle progression via the downregulation of the expression of cyclins in *Atg7-/-* mice. These results indicate that hepatic tumors of *Atg7-/-/Hmgb1-/-* are less proliferative than the tumors in *Atg7-/-* mice. Thus HMGB1 had an impact on cell proliferation in the autophagy-deficient liver.

We then examined the phosphatidylinositol 3-kinase(PI3K)/AKT signaling pathway that regulates various cellular responses in HCC proliferation and survival ^27, 28^. Intriguingly, immunoblot analysis showed that phospho-AKT was detected at higher levels in *Atg7-/-/Hmgb1-/-* livers compared to *Atg7-/-* livers regardless the sample type (**Supplementary Figure S7A**). The expression level of phospho-glycogen synthase kinase 3 (GSK3)β, one of the downstream target of AKT was also markedly increased in the *Atg7-/-/Hmgb1-/-* livers (**Supplementary Figure S7A**). However, the protein levels of total and phospho-PDK, an upstream regulator of AKT were comparable between the two genotypes (**Supplementary Figure S7A**). In addition, we found an increased level of phospho-JNK in *Atg7-/-/Hmgb1-/-* livers as compared to that in *Atg7-/-* livers regardless the sample type (**Supplementary Figure S7B**). JNK can be a dominant effector of mitogen-activated protein kinase in the liver ^29^. JNK catalyzes the phosphorylation of numerous substrate proteins including the c-Jun transcription factor to regulate the gene expression. However, the protein expression level of phospho-c-Jun and total-c-Jun were comparable between the two genotypes (**Supplementary Figure S7B**). The mammalian target of rapamycin complex 1 (mTORC1) signaling pathway, the mitogen-activated protein kinase(MAPK)/ERK signaling pathway, and the Janus activated kinase/Signal Transducer and Activator of Transcription Family or transcription factors (JAK/STAT3) signaling pathway, all have been associated with cell growth ^30–33^. However, we did not detect significant differences in the activation of these pathways between *Atg7-/-* mice and *Atg7-/-/Hmgb1-/-* mice (**Supplementary Figure S7C-E**). Taken together while the reason for the paradoxical elevation of AKT and JNK phosphorylation in *Atg7-/-/Hmgb1-/-* livers is not clear these events do not seem to be tumor specific and may not be related to the reduced proliferation status of tumors from in these livers. Alternativley, it is notable that the hepatocytes could offer a very different cellular context in which the conventional oncogenes or tumor suppressor genes can act in opposite ways ^34, 35^.

### 7. RAGE deletion impairs proliferation and retards liver tumor development

HMGB1 is a non-histone nuclear protein that facilitates the binding of regulatory proteins to DNA and typically enhances transcriptional activation^10^. When released extracellularly, HMGB1 can binds to one of its receptors, such as RAGE or TLR4 ^36^. In our previous study, *Atg7-/-* mice develop hepatic tumors at 9-month old in the liver, which could be inhbited by the deletion of either *Hmgb1* or *Rage* ^2^. While *Atg7-/-/Hmgb1-/-* at the age of 12-month old were still largely devoid of tumors in the liver ^2^, we now found that 12-month old *Atg7-/-/Rage-/-* mice developed a significant presence of tumors albeit at a level slightly lower than that in the *Atg7-/-* livers (**Figure 6A**). However, hepatic tumors developed in the 12-month old *Atg7-/-/Rage-/-* mice were significantly smaller in size compared to those in the *Atg7-/-* mice (**Figure 6B**). The data indicate that deletion of *Rage* still delayed the development of tumors in older *Atg7-/-* mice although in a less prominent manner than *Hmgb1* deletion. Notably, the number of PCNA-positive cells and the expression of cyclin D1 were also remarkably decreased in *Atg7-/-/Rage-/-* livers compared to that in the *Atg7-/-* livers(**Figure 6C-D**). These data suggest that the loss of RAGE in autophagy-deficient livers reduced tumor cell proliferation and tumor expansion in the liver. HMGB1 interaction with the RAGE receptor can thus mediate a significant level of cell proliferation and tumor development in the autophagy-deficient liver.

**Figure 6.**
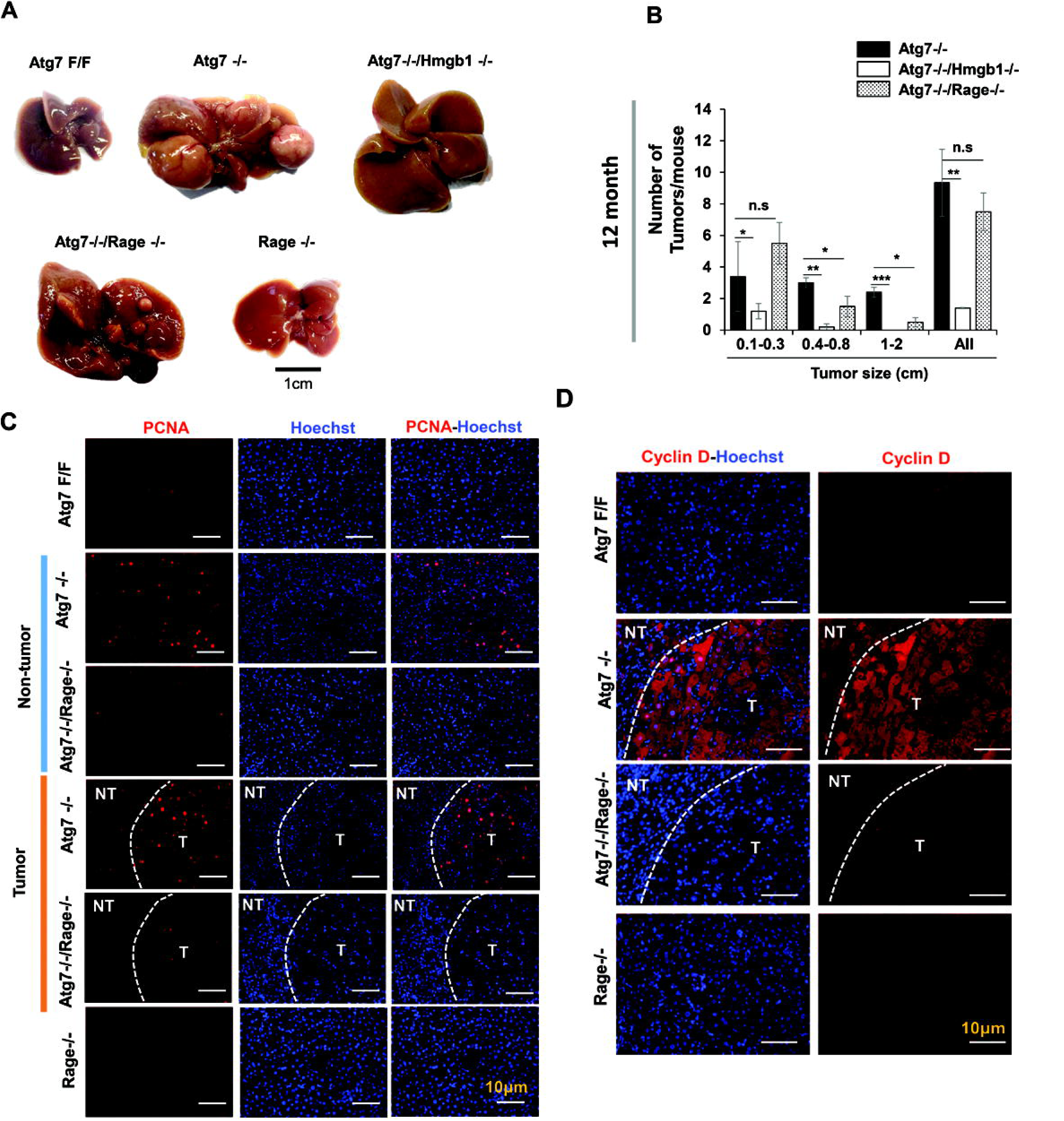
Genetic loss of *Rage* inhibits tumorigenesis in autophagy-deficient livers. (**A**) Gross images of representative livers of 12-month old *Atg7-/-*, *Atg7-/-/Hmgb1-/-*, *Atg7-/-/Rage-/-*, and *Rage-/-* mice. (**B**) Average number and size distribution of the tumors observed in the livers of 12-month old mice of different genotypes. (**C-D**) Liver sections from 12-month old mice of different genotypes were immunostained with anti-PCNA(C), or anti-Cyclin D (D). White dotted lines indicate the tumor border. NT. non-tumor, T, tumor. Data are reported as mean± SE,* *P*<0.05, ** *P*<0.01, *** *P*<0.001, n.s., no significance; n=3 mice per group. Size information of the tumor from *Atg7-/-/Hmgb1-/- livers* is derived from what we has previously reported^2^.

To determine whether HMGB1 released by the autophagy-deficient hepatocytes or hepatic tumor cells could act as an autocrine fashion to promote cellular proliferation, we examined whether hepatoctyes could express RAGE. Immunofluorescence staining was performed in frozen tissue from *Atg7 F/F* and *Atg7-/-* liver. We found that RAGE was almost exclusively expressed by cells other than hepatocytes based on cell morphology, but it was detected on the cell surface, consistent with its being a receptor molecule (**Figure 7A**). To examine which non-hepatoctyes expressed RAGE, double immunofluorescence staining for RAGE, together wtih CK19 or SOX9(for ductular cells), F4/80 (for Kupffer cells), or Desmin (for stellate cells), was conducted. Colocalization of RAGE was evident in CK19 or SOX9-positive ductular cells and F4/80-positive Kupffer cells, but not on the Desmin-positive stellate cells in *Atg7-/-* liver (**Figure 7B**).

**Figure 7.**
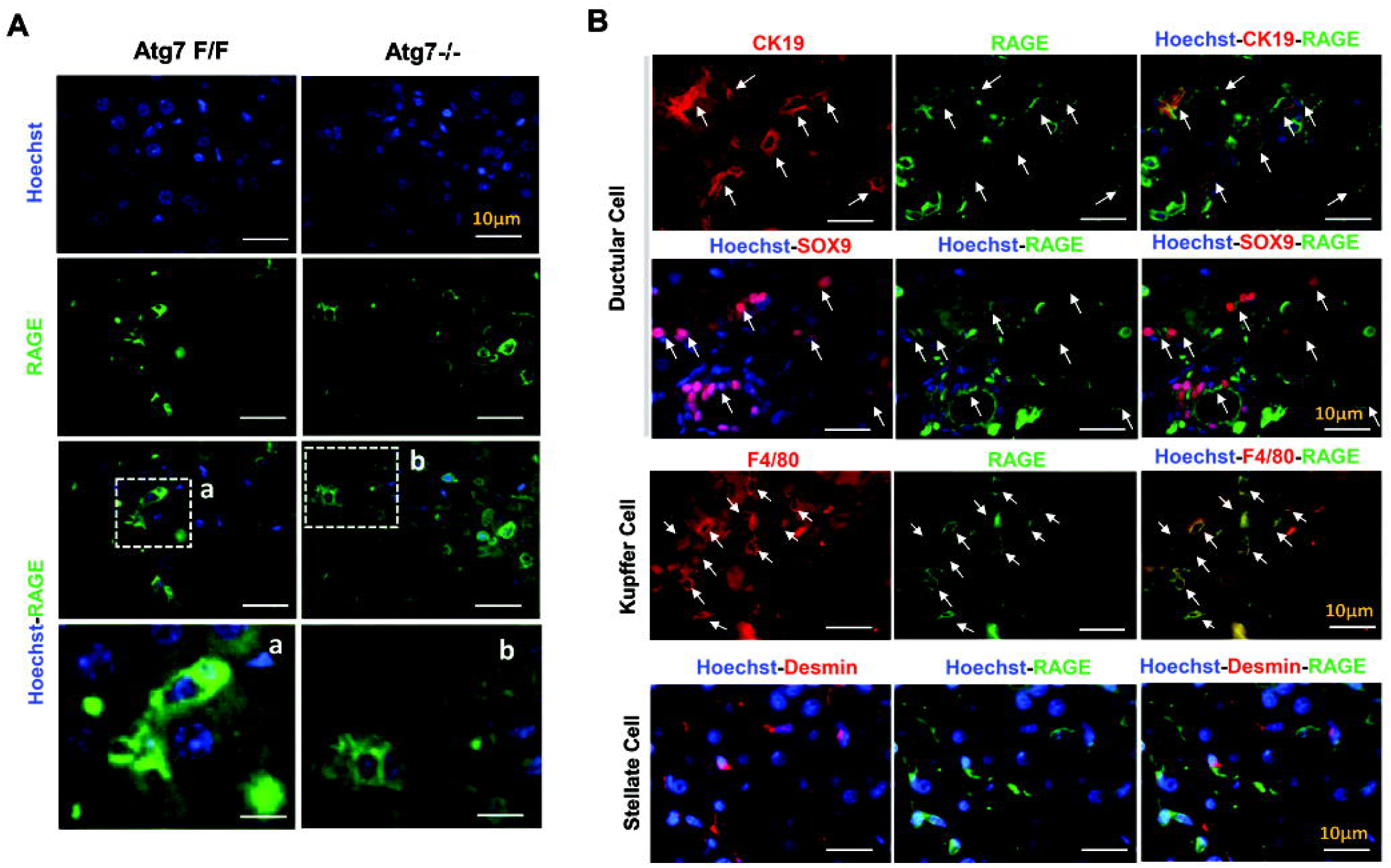
RAGE is expressed by ductular cells and Kupffer’s cells but not by hepatocytes or stellate cells. (**A**) Immunofluorescence staining for RAGE antigen in the livers of 9-week old mice of *Atg7F/F* and *Atg7-/-* genotype. Framed ares are enlarged and shown in separate panels (a,b). (**B**) Liver sections from 9-week old *Atg7-/-* mice were coimmunostained with anti-RAGE, together with anti-CK19 or SOX9 or F4/80 or Desmin. White arrows indcate cells with colocalized signals.

These findings indicate that RAGE was expressed on ductular cells and Kupffer cells but not on hepatocytes nor stellate cells. Futhermore, these observations suggest that unlike the possible direct effect of HMGB1 on the expansion of CK19-positive or SOX9-posotivie ductual cells ^2^, the tumor-promoting effect of HMGB1 may not be mediated by a direct effect on the autophagy-deficient hepatocytes, but possibly by an indirect effect through other RAGE-expressing cells, such as the Kupffer’s cells, which could then alter the microenvironment that facilitate tumor development.

### 8. RNA Sequencing revealed key molecular differences between tumors from *Atg7-/-* mice and from *Atg7-/-/Hmgb1-/-* mice

Since the effect of HMGB1 in promoting tumor development may be mediated by an altered microenvironment, there could be multiple alterations in tumor behaviors affected by this process. We sought to investigate the transcriptomic profile of the tumor to better understand the impact of HMGB1 on tumor development in autophagy-deficient livers. We chose to perform RNA sequencing on tumor tissues obtained from *Atg7-/-* and *Atg7-/-/Hmgb1-/-* mice at the age of 15 months old, when the tumor number and size were comparable in these mice.

The principal component analysis (PCA) on the RNAseq data indicated different transcriptomic profiles in the tumor tissues of 15-month old *Atg7-/-* and *Atg7-/-/Hmgb1-/-* mice when compared with the non-tumor tissues (**Figure 8A**), The six non-tumor samples from the two strains of mice were close to each other. In addition, two out of the three tumor samples from *Atg7-/-/Hmgb1-/-* livers were also close to the non-tumor samples whereas tumor samples from *Atg7-/-* mice were separated the farthest from the rest of the samples. PCA thus suggests that tumors from the two strains of mice were quite different with those from *Atg7-/-/Hmgb1-/-* livers more similar to the non-tumor tissues in their transcriptomic profiles.

**Figure 8.**
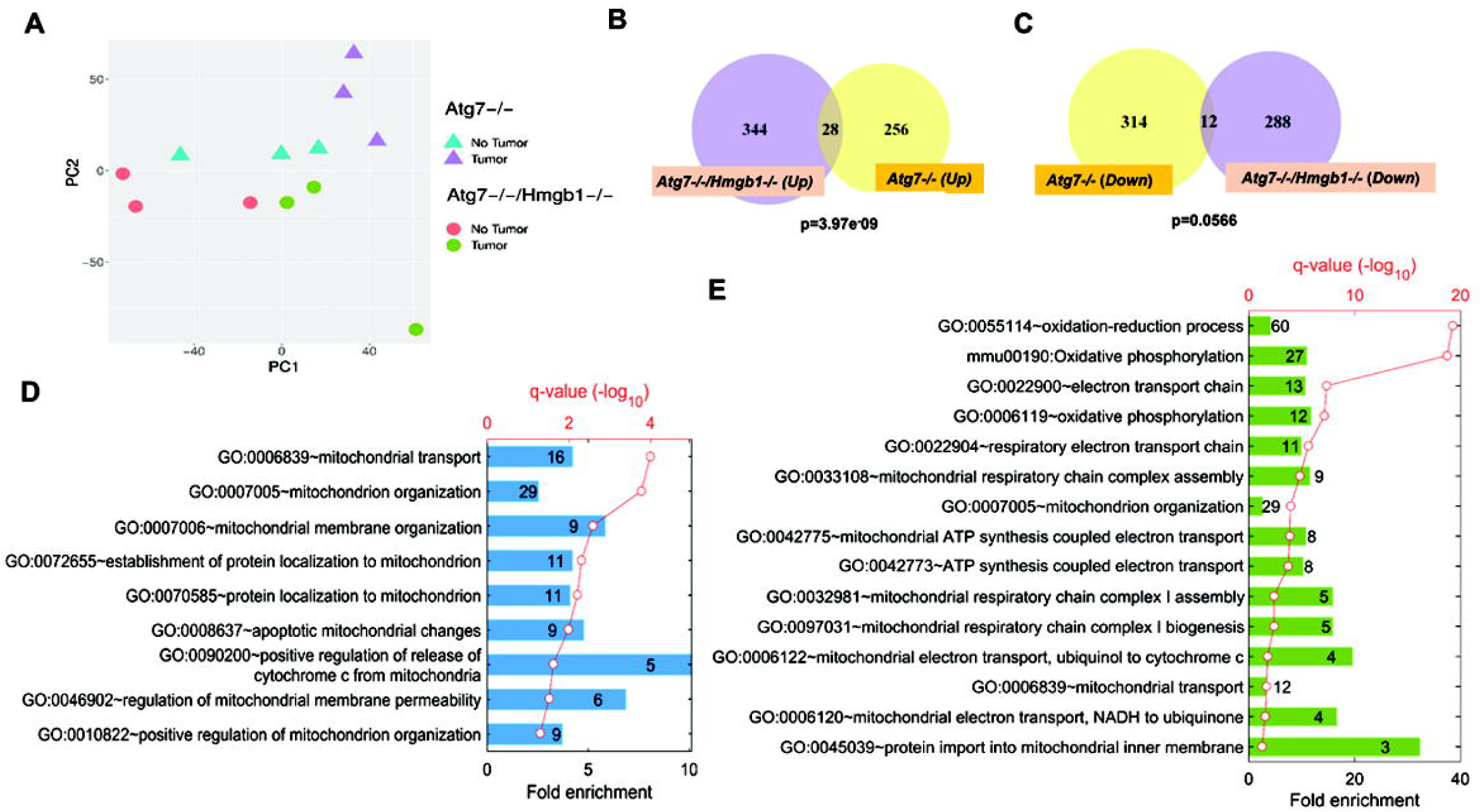
RNAseq analysis indicates transcriptomic diffrences in the hepatic tumors of *Atg7-/-* mice and *Atg7-/-/Hmgb1-/-* mice. **(A**). PCA of transcriptomic data based on 12 RNA-seq samples under the four indicated combinations of genotyes and tissue types. (**B-C**). Numbers of DEGs that are significantly up-regulated (B) or down-regulated (C) (p<0.01) in the tumor samples of *Atg7-/-/Hmgb1-/-* and/or *Atg7-/-* mice. The p-values are indicated for the overlap between the two groups of upregulated or downregulated DEGs, respectively. (**D**). GO biological processes significantly over-represented in the non-overlapped 256 DEGs uniquely elevated in the tumor samples of the *Atg7-/-* mice. (**E**). GO biological processes and KEGG pathways significantly enriched in the non-overlapped 288 DEGs uniquely repressed in the tumor sampels of the *Atg7-/-/Hmgb1-/-* mice. For D and E, the heights of bars indicate the fold enrichment compared to random selection, whereas the red dots represent the statistical significance, p-value after FDR-adjusted multiple test correction. The numbers in the bars represent the numbers of DEGs in the particular group which are associated with corresponding GO terms.

Differential expression analysis showed that 284 and 372 differentially expressed genes (DEGs) were upregulated in tumors of *Atg7-/-* and *Atg7-/-/Hmgb1-/-* livers, respectively, whereas 326 and 300 genes were downregulated in tumors of these livers, respectively (**Figure 8B-C**). A complete list of these DEGs can be found in **Supplementary Tables S1-S5**. We then focused on discovering unique molecular features in the tumous associated with the presence and absence of HMGB1. When comparing the DEGs between *Atg7-/-* and *Atg7-/-/Hmgb1-/-*, a small number of up-regulated (28, **Figure 8B**) or down-regulated (12, **Figure 8C**) DRGs were found in tumor tissues of both *Atg7-/-* and *Atg7-/-/Hmgb1-/-* livers. The larger portions of DEGs were, however, unique in *Atg7-/-* and in *Atg7-/-/Hmgb1-/-* tumors, supporting that the tumors were different in the presence or absence of HMGB1.

To understand the molecular features of these differences, we determined the Gene Ontology (GO) terms and KEGG pathways that were significantly enriched in the unique DEG sets. We found that many biological processes, particularly those associated with mitochondrial structrures or functions were significantly over-represented by the uniquely up-regulated DEGs in *Atg7-/-* tumors (**Figure 8D**). Notably, DEGs down-regulated uniquely in *Atg7-/-/Hmgb1-/-* tumors were also enriched for those involved in the mitochondrial structures or functions, (**Figure 8F**). Many genes related to mitochondrial oxidative phosphrlation(OXPHOS) or electron transport chain (ETC) process were significantly downregulated in the tumors of *Atg7-/- Hmgb1-/-* liver. Particulalry, the genes involved in the assemby or biogenesis of respiratory complex I (NADH dehydrogenase complex) and complex III(Ubiquinol to Cytochrome c electron transporter) were significantly downregulated. These observations suggested that a major component of the tumor-promoting effects of HMGB1 could be related to mitochondrial function and activity, which may impact the celluar bioenergetics and hence tumor growth in autophagy-deficient liver.

## Discussion

Autophagy is important for liver homeostasis and tumor surveillance. Deficiency of hepatic autophagy, such as that caused by the liver-specific deletion of *Atg5* or *Atg7*, leads to tumor development in aged mice ^1–4^. On the other hand, autophagy function is required for the aggressive growth of tumors. The mechanism that sustains the growth of autophagy-deficient tumors is not known. In this study, we examined the cellular and molecular nature of hepatic tumors in the autophagy-deficient liver. Our findings support the following conclusions: 1) The adenoma originates from the autophagy-deficient hepatocytes; 2) Hepatocyte-derived HMGB1 stimulates tumor cell proliferation; 3) HMGB1 mediates the proliferative signal at least in part via RAGE in a paracrine mode; and 4) Tumors developed in the presence or absence of HMGB1 have significantly different transcriptomic profiles and mitochondria function could be an important mechanistic linker to tumor promotion.

### 1. HMGB1 may act in a paracrine model to stimulate tumor growth

Hepatic tumors were histologically consistent with hepatocellular adenoma where the benign tumor cells arrange in regular plates, usually one or two cells in thickness ^3^. Here we further showed that tumor cells were originated from the autophagy-deficient hepatocytes.

The composition of the tumor appears to be different from the non-tumor liver tissue. In comparison to non-tumor tissues where different hepatic cells including hepatocytes, inflammatory cells, fibrotic cells, and ductular cells coexist, the tumor tissue consists of mainly the tumor cells (HNF4α positive), and some macrophages (**Figure 1–3**)(**Supplementary Table S6**). Other nonparenchymal cells are found only outside the tumor region. Fibrotic cells and ductular cells seem to be responsible for the formation of a fibrous capsule that demarcates the tumor from the non-tumor tissue. How the autophagy-deficient hepatocytes form the adenomatous nodule, excluding the fibrotic cells and ductular cells but retaining some macrophages, is intriguing. But macrophages could belong to those known as tumor-associated macrophages(TAM) and may enter into the tumor tissue via tumor blood vessels ^37^.

HMGB1 is known to promote tumor development ^2, 38, 39^. However, how HMGB1 does this, in particular for the autophagy-deficient heaptic tumors, is not clear. HMGB1 has been shown to be important for expansion of ductular cells ^2, 38^, immune cells recruitment ^25^ and activation of fibrotic cells ^24^. All these cellular events could favor tumorigenesis. Previously, we showed that non-tumorigenic autophagy-deficient hepatocytes can release HMGB1 ^2^. Here our findings indicate that HMGB1 could be also released by the autophagy-deficient hepatic tumor cells. Thus HMGB1 may act through an autocrine or paracrine mode to promote tumor growth.

We found that the proliferative effect of HMGB1 could be mediated via the RAGE receptor as deletion of *Rage* also reduces tumor cell proliferation and delay tumor development (**Figure 6**). We found that RAGE receptor was not expressed by hepatocytes and stellate cells at the detectable level by immunostaining. But it could be readily detected on the surface of Kupffer’s cells and HPCs ^2 40^. Hence. the effect of HMGB1 in cell proliferation could be mediated by a paracrine manner, although the autocrime mode could be not be completedly excluded (**Supplementary Figure S8**). In the paracrine mode, HMGB1 release by the tumorigenic and non-tumorigenic autophagy-deficient hepatocytes could activate macrophages or ductular cells, which then releases different cytokine factors that ultimately affect hepatocytes growth and proliferation. From this aspect, interaction between tumorigenic hepatocytes and TAM could generate an intratumoral microenvironment favoring cell growth and proliferation. However, it seems that some of the well-defined proinflammatory cytokines such as TNFα, IL-1β, and IL-6 may not play the role as the expression of these cytokines are remarkably downregulated in tumor tissues of *Atg7-/-* mice (**Supplementary Figure S4**). On the other hand, the RAGE-positive peri-tumoral ductular cells could possibly communicate with hepatocytes via cytokines such as angiogenic factors ANGPT2, PDGFb to promote protumorigenic activities, such as angiogenesis and invasiveness while inhibiting tumor infiltration into normal tissue. It is also possible that the protumorigenic factors from TAM and/or ductular cells could be mediated by extracellular vesicles, microRNAs and other cellular factors ^41^.

The *Rage* deletion could not fully protect the *Atg7-/-* mice from tumor development at age of 12-month oold although the tumor size still appearred significantly smaller (**Figure 6A-B**). This is in contrast to *Atg7-/-/Hmgb1 -/-* mice where *Hmgb1* deletion could significantly inhibit tumor development in *Atg7-/-* mice even at the 12-month time point (**Figure 6A-B**)^2^. It is possible that HMGB1 affect the tumorigenesis process not only via RAGE but also through other receptors such as TLR4 ^36^. Future studies can assess the potential role of TLR4 in this process.

### 2. The impact of HMGB1 on tumor cells can be broad

Deletion of *Hmgb1* or *Rage* led to a signficant reduction in the proliferative capability of autophagy-deficient hepatocytes and tumors as demonstrated by the expression of PCNA, Ki67 and Cyclin D1. Thus the pro-proliferative effect by HMGB1 confers a geneally stronger capability of proliferation to autophagy-deficient hepaotcytes, which would be benefical to the growth of tumors that are derived from these cells.

However, RNAseq analysis indicates that there are much more unique changes in the molecular composition of the tumors caused by HMGB1. The enrichment of certain gene expression related to mitochondrial structure and function in the presence of HMGB1 and lack of such enrichment in the absence of HMGB1 are quite significant. *Hmgb1* deletion appears to suppress the mitochondrial ETC in tumors of autophagy-deficient livers. Whether and how downregulation of genes of mitochondrial ETC may suppress cell proliferation in *Atg7-/-Hmgb1-/-* tumors is unclear. But it is well known that mitochondrial ETC enables many metabolic processes and is a major sources of ATP and building blocks for the proliferation of tumor cells. As a consequence of ETC dysfunction, cell proliferation could be impaired due to bioenergetics deficit. This notion is supported by the observation where pharmacological or genetic inhibition of ETC caused impaired cell proliferation of cells in vitro ^42, 43^. Interestingly, a recent study suggest that ETC enables aspartate biosynthesis, a key proteogenic amino acid that is also a precursor in purine and pyrimide synthesis and is required for tumor cells growth and survival ^44, 45^. Thus tumors of *Atg7-/-Hmgb1-/-* liver may have defective ETC that could impair cell proliferation by limiting an intracellular aspartate level besides causing bioenergetics deficits. Many metabolic pathways including glycolysis, the TCA cycle, and β-oxidation produce the electron donors that fuel the ETC. Hence, impairment or downregulation of ETC could limit the regeneration of reducing equivalents, such as NAD+, which in turn suppresses glycolysis or the TCA cycle. Future studies should address these possibilities for the understanding of how HMGB1 sustains the growth of autoaphgy-deficient hepatic tumors.

In conclusion, our findings demonstrate that hepatic adenoma originates from the autophagy-deficient hepatocytes that release HMGB1. HMGB1, in turn, can stimulate hepatocyte proliferation and hepatic tumorigenesis via RAGE in the autophagy-deficient liver. The effect of HMGB1 on tumor cells are broad as revealed by transcriptomic analysis, which offers mechanistic clues for future studies.

## Materials and Methods

### Animal experiments

*Atg7F/F*, *Atg7-/-*, *Atg7-/-/Hmgb11-/-*, *Atg7-/-/Rage-/-*, *Hmgb1-/-,* and *Rage-/-* mice were used in this study. *Atg7F/F* was obtained from Dr. Komatsu Masaaki (Nigata University, Japan). These mice were backcrossed with C57BL/6J for another 10 generations as described previously^2, 20^. Albumin-Cre mice were obtained from the Jackson Laboratory(Bar Harbor, ME). *Hmgb1 F/F* and *Rage* mice were as described ^2^. Hepatic *Atg7-/-* mice were further crossed with *Hmgb1 F/F* or *Rage* to generate *Atg7-/-/Hmgb1-/-* or *Atg7-/-/Rage-/-* mice as previously described^2^. Both male and female mice were used in the study. All animals received humane care, and all procedures were approved by the Institutional Animal Care and use Committee(IACUC) of the Indiana University.

### Tumor sample collection

The whole liver was carefully removed from the euthanized animals, washed, and placed in cold PBS. The number of tumor nodules on the liver surface was counted for all the liver lobes. Tumor nodules with >2mm in diameter were carefully removed and examined as tumor tissue. Tissue without visible tumor nodules were sampled as non-tumor tissues. All tissues were collected in separate tubes and stored at -80°C for future studies. Liver tissues containing the tumor nodule and the surrounding non-tumor tissue were excised and fixed in 10% neutral formalin or buffered with 4% PFA overnight for paraffin-embedding or for OCT embedding. The tissue section was prepared from the frozen or paraffin blocks for general histology, immunostaining, and immunohistochemistry analysis.

### General histological and immunological analysis

General histology was examined on paraffin-embedded sections stained with hematoxylin and eosin (H-E). Liver fibrosis was determined by Sirius Red staining or Masson’s Trichome staining. For immunostaining, liver sections were subjected to heat-induced antigen retrieval using citrate buffer (pH 6.0) followed by permeabilization and blockage with 10% goat or donkey serum in PBS containing 0.5% triton-X for 1 hour. Sections were incubated overnight at 4°C with primary antibody diluted in PBS. Primary antibodies used in this study are listed in **Supplementary Table S8**. Sections were then incubated with Alexa-488 or Cy3-conjugated secondary antibodies. Images were obtained using Nikon Eclipse TE 200 epi-immunofluorescence microscope. Hoechst 33342 was used for nucleus staining. Images were analyzed using NIS-element AR3.2 software.

Immunoblot analysis was performed as described previouosly^2, 20^ using primary antibodies and respective secondary antibodies conjugated with horseradish peroxidase as listed in **Supplementary Table S8**. The respective protein bands were visualized using the immunobilion chemiluminescence system (Millipore, MA). The densitometry analysis of immunoblotting images was performed using Quantity One Software (Bio-rad). Densitometry values were normalized to the loading control (GAPDH) and then converted to units relative to the untreated control.

### Total RNA isolation, reverse transcription, and quantitative real-time PCR analysis

Total RNA was isolated from liver tissues using a GeneTET RNA Purification Kit (Thermo Fisher Scientific) according to the manufacturer’s protocol. cDNA was synthesized using an M-MLV Reverse Transcriptase Enzyme System (Life Technologies, Thermo Fisher Scientific) and OligodT primers. The resulting cDNA products were subjected to qPCR reaction using SYBR Green Master Mixes. qPCR was performed on a Quanta studio 3 PCR machine (Life Technologies–Applied Biosystems, Thermo Fisher Scientific). The threshold crossing value(Ct) was determined for each transcript and then normalized to that of the internal gene transcript(β-actin). Fold change values were then calculated using the 2^−ΔΔCt^ method. Genes-specific primers were designed using Integrated DNA Technologies (IDT) PrimerQuest software. Sequences of the forward and reverse primers are listed in **Supplementary Table S7**.

### RNA-sequencing and bioinformatics analysis

RNA was isolated as described above. RNAseq was performed by The Center for Medical Genomics facility at Indiana University. The integrity of RNA was determined using an Agilent Bioanalyzer 2100 (Agilent Technologies;Santa Clara, CA). Extracted RNA was processed for rRNA removal using the Epicenter rRNA depletion kit according to the manufacturer’s instructions. rRNA-depleted RNA was subsequently used to generate paired-end sequencing libraries using the Illumina RNA TruSeq Library Kit according to the manufacturer’s instruction. RNAseq was performed using Illumina HiSeq 4000 (Illumina, San Diego, CA). For bioinformatics analysis, we first used FastQC (http://www.bioinformatics.babraham.ac.uk/projects/fastqc) to examine RNA-seq quality. Then all high-quality sequences were mapped to the mouse genome (mm10, UCSC Genome Browser, https://genome.ucsc.edu/) with the STAR, an RNA-seq aligner^46^. The featureCounts was adopted to assign uniquely mapped reads to genes according to UCSC refGene (mm10) ^47^. Those low-expressed genes were not further analyzed if their raw counts were less than 10 in more than three samples for each pairwise comparison. The gene expression was normalized cross all samples based on trimmed mean of M (TMM) values implemented in EdgeR ^48^, followed by differential expression analysis given comparisons between non-tumor and tumor tissues, in either single knockout or double knockout mice. Genes with p values less than 0.01 after multiple-test false discovery rate (FDR) correction were determined as differentially expressed genes (DEGs) for specific comparisons. The gene ontology (GO) and KEGG pathways significantly enriched in DEGs were identified by DAVID functional annotation analysis tools^49^.

### Statistical Analysis

Statistical analyses were performed with Sigma Plot. All experimental data were expressed as Mean±SE. Student t-test was performed to compare values from two groups. To compare values obtained from three or more groups, one way ANOVA analysis with the appropriate post-hoc analysis was used. Statistical significance was taken at the level of P< 0.05.

## Supporting information

Supplementary Table S1-Gene Lists-Overlapped

Supplementary Table S2-Gene Lists- Atg7KO only upregulated

Supplementary Table S3-Gene Lists- Atg7HMGB1DKO only upregulated

Supplementary Table S4-Gene Lists- Atg7KO only downregulated

Supplementary Table S5-Gene Lists- Atg7HMGB1DKO only downregulated

Supplementary Table S7-List of Primers used for qPCR

Supplementary Table S8-List of Antibodies used for immunostaining and western blot

## Acknowledgment

The CK19 monoclonal antibodies were obtained from the Developmental Studies Hybridoma Bank, created by the National Institute of Child Health and Human Development (NICHD), NIH, and maintained at the Department of Biology of The University of Iowa (Iowa City, Iowa, USA). This work was in part supported by NIH/NIDDK grants R01 DK116605 (XMY).

## Authors contributions

BK designed project and directed study, analyzed data and wrote the manuscript. BK, XC conducted experiments, acquired data, and analyzed data. HH, GL, JW helped in gene expression analysis. ZD gave critical discussions. XMY designed research experiments, analyzed data, and edited the manuscript.

