## Supplementary Figures and Figure legends for "The HMGB1-RAGE axis modulates the growth of autophagy-deficient hepatic tumors"

1) **Supplementary Figure legends**

2) **Supplementary Figures S1-S8 and Supplementary Table S6**

Khambu B et al 2019

#### Supplementary Figure legends

**Supplementary Figure S1.** SOX9 positive hepatic progenitor cells are localized outside the tumor regions. Liver sections from 12-month old mice of *Atg7-/-* genotype were co-immunostained with anti-SQSTM1 and anti-SOX9. Images were taken around the peri-tumor and tumor region (**A**). Additional higher magnification images in the peri-tumor regions (**B**) were taken in order to show the SQSTM1 staining in SOX9 positive hepatic progenitor cells. The framed area is enlarged and shown in separate panels. White dotted lines indicate the tumor border. NT, non-tumor liver, B, peri-tumor, T, tumor.

**Supplementary Figure S2**: Elevation of Cancer stem cell-associated genes in non-tumor and tumor samples of autophagy-deficient livers. The hepatic mRNA expression level of CSCs marker genes (*Cd133/Prom1, Cd200/Ox-2,Cd34, Cd44, Ly6a/Sca-1, Ly6d, Cd24A/Has, Cd90/Thy1)* (**A**), and Stemness genes (*Oct4, Nanog, Klf4,* and *Sox2*) (**B**) in 15-month old *Atg7F/F,* and *Atg7-/-* mice were determined by real-time PCR. NT. non-tumor, T, tumor. Data are reported as mean± SE,* $P<0.05$, *** $P<0.00$, n.s.: no significance; n=3 mice per group.

**Supplementary Figure S3.** Angiogenic factors are altered in the tumor-bearing autophagy-deficient livers**.** The hepatic mRNA expression level of angiogenic factors (*Angpt2, Pdfgb Vegfra,* and *Angpt1)* in 15-month old *Atg7F/F* and *Atg7-/-* mice were determined by real-time PCR. NT, non-tumor, T, tumor. Data are reported as mean± SE,* $P<0.05$, ** $P<0.01$, n.s., no significance; n=3 mice per group.

**Supplementary Figure S4.** Expressional analysis of proinflammatory cytokine genes in tumor and non-tumor samples of autophagy-deficient livers. The hepatic mRNA expression level of inflammatory cytokines (*TNFα, IL-6, IL-1β* and, *IL-17)* in 15-month old *Atg7F/F* and *Atg7-/-* mice were determined by real-time PCR. NT, non-tumor, T, tumor. Data are reported as mean± SE,* $P<0.05$, n.s., no significance; n=3 mice per group.

**Supplementary Figure S5.** The autophagy-deficient tumors are proliferative. Liver sections from 12-month old mice of *Atg7-/-* genotype were subjected to immunohistochemistry for Ki67

(**A**)  (original magnification, X200). Dotted lines indicate the tumor border.  (**B**) Enlarged images of Region 1 (peri-tumor), Region 3 (tumor) and Region 7 (non-tumor) are shown in separate panels. Red arrow indicated Ki67 positive proliferating hepatocytes. NT, non-tumor, T, tumor.

**Supplementary Figure S6.** Cyclin D expression in autophagy-deficient livers. Livers of 15-month old mice of *Atg7-/-* and *Atg7-/-/Hmgb1-/-* genotypes were immunostained with anti-Cyclin D. Several images were taken focusing in the tumor region. The framed area is enlarged and shown in separate panels. White dotted lines indicate the tumor border.

**Supplementary Figure S7.** Loss of *Hmgb1* activates AKT and JNK signaling but does not affect mTORC1, MAPK/ERK and STAT signaling in the autophagy-deficient livers. (**A-B**) Immunoblot analysis of  AKT pathway (A)  and  JNK signaling pathway (B) related proteins in the tumor or non-tumor sample of 15-month old *Atg7-/-* and, *Atg7-/-/Hmgb1-/-* mice. (**C-E**) Immunoblot analysis of  mTORC1 signaling (C), MAPK/ERK signaling (D), and STAT signaling (E) related proteins in the tumor or non-tumor samples of 15-month old *Atg7-/-* and, *Atg7-/-/Hmgb1-/-* mice. NT, non-tumor, T, tumor.

**Supplementary Figure S8.** Schematic model for the role of HMGB1 in tumor development in the autophagy-deficient liver.  HMGB1 is released from autophagy-deficient hepatocytes via the NRF2-inflammasome pathway.  Deletion of RAGE, an HMGB1 receptor, mimicked the effect of HMGB1 deletion in delaying tumor development, suggesting that HMGB1 affects tumor development via its released form, but not its DNA-binding form.  That HMGB1 may act on hepatocytes in an autocrine fashion could not be completely excluded although hepatocytes do not seem to express a detectable level of RAGE.  Released HMGB1 could thus have paracrine effects on target cells that express RAGE and may affect tumor development by altering the microenvironment.

**Supplementary Tables**

1. **Supplementary Table S1.** List of overlapped genes that are upregulated or downregulated in tumors of both *Atg7-/-* and *Atg7-/-/Hmgb1-/-* liver.
2. **Supplementary Table S2.** List of upregulated genes in tumors of *Atg7-/-* liver.
3. **Supplementary Table S3.** List of upregulated genes in tumors of *Atg7-/-/Hmgb1-/-* liver.
4. **Supplementary Table S4.** List of downregulated genes in tumors of *Atg7-/-* liver.
5. **Supplementary Table S5.** List of downregulated genes in tumors of *Atg7-/-/Hmgb1-/-* liver.
6. **Supplementary Table S6.** Summary of distribution of hepatic cells in non-tumor, peri-tumor and tumor tissues of the autophagy-deficient liver.
7. **Supplementary Table S7.** List of Primers used for qPCR.
8. **Supplementary Table S8.** List of Antibodies used for immunostaining and western blot.

### Supplementary Figure S1

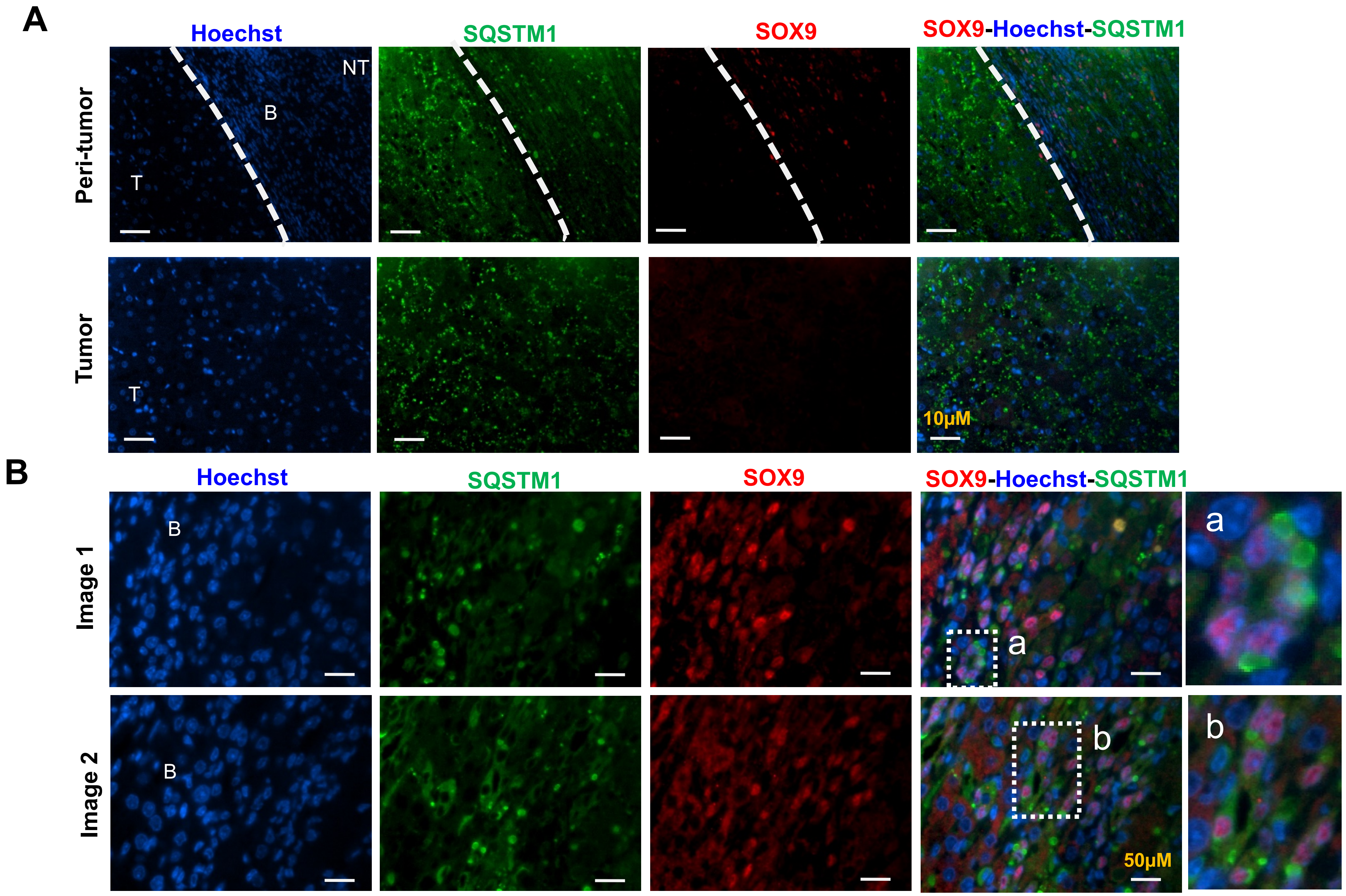

#### Supplementary Figure S2

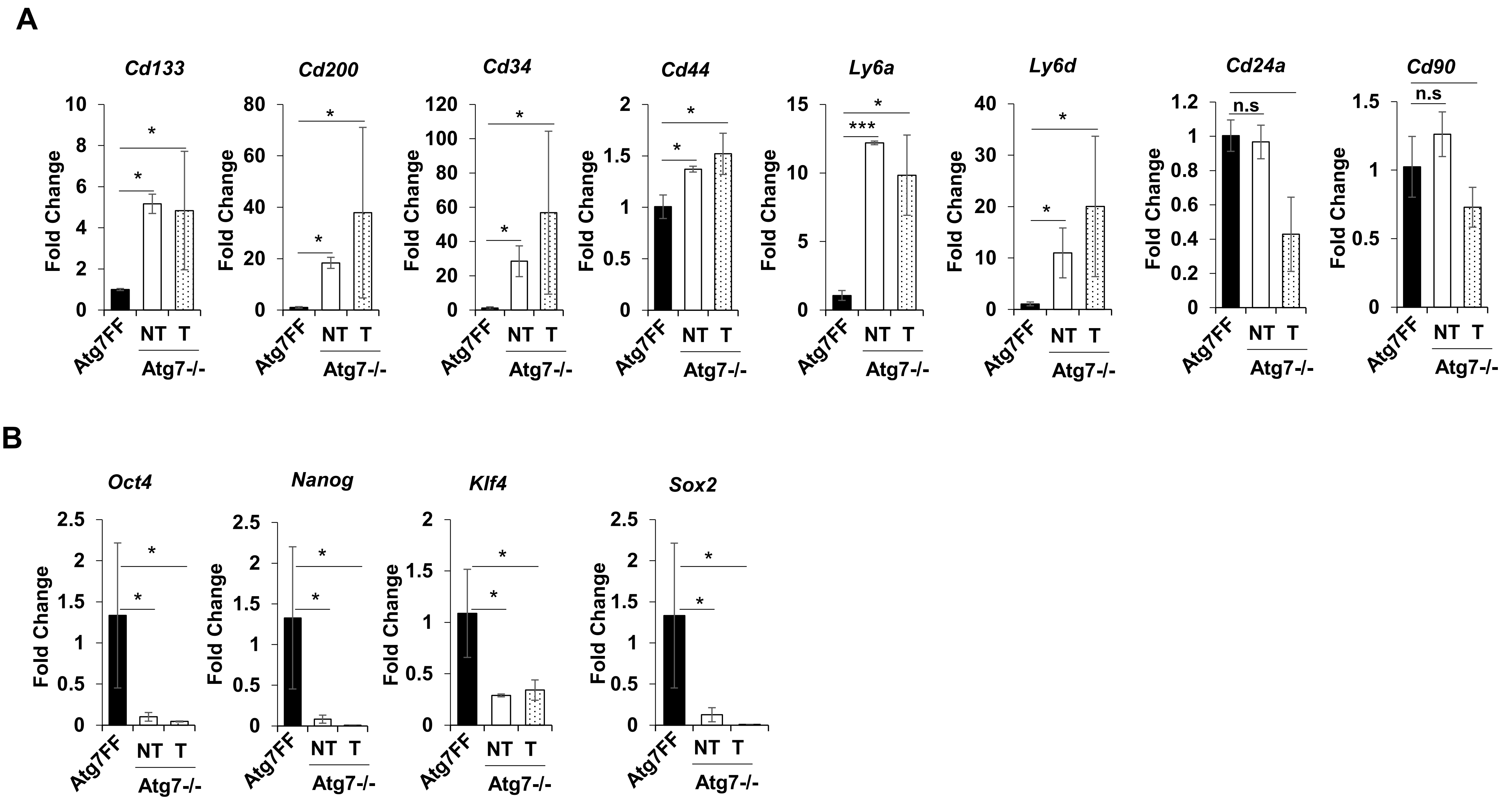

### Supplementary Figure S3

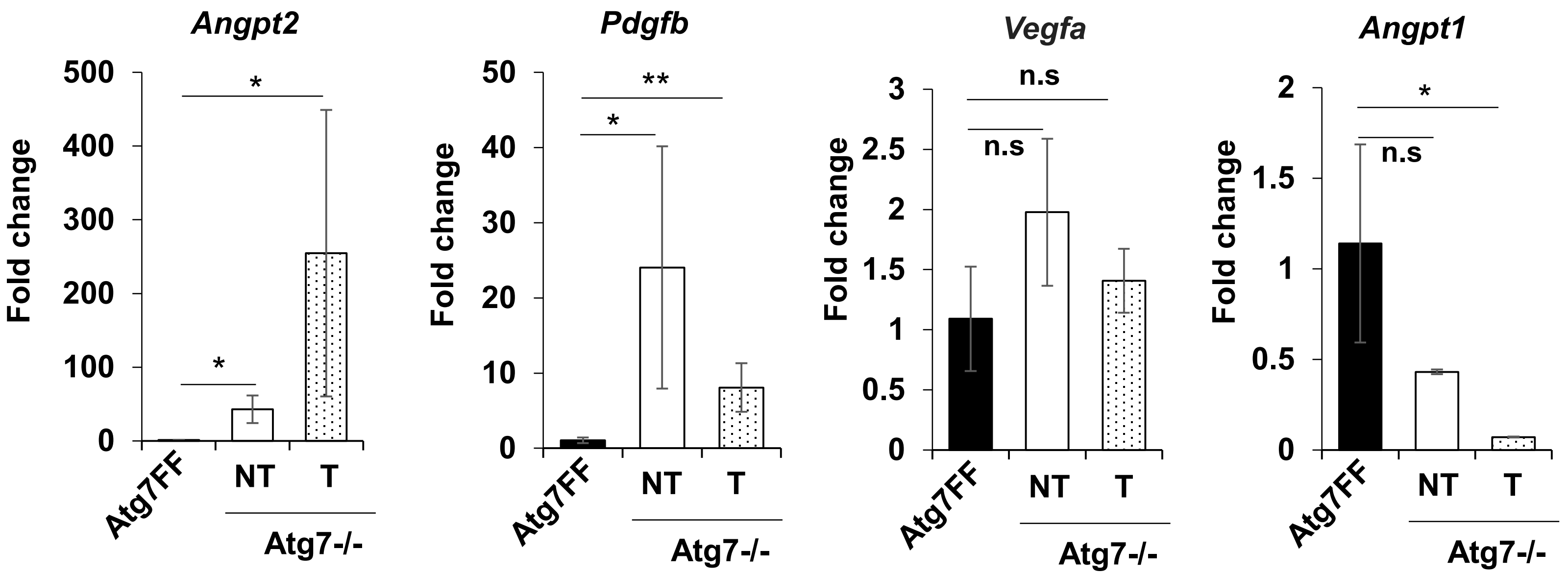

#### Supplementary Figure S4

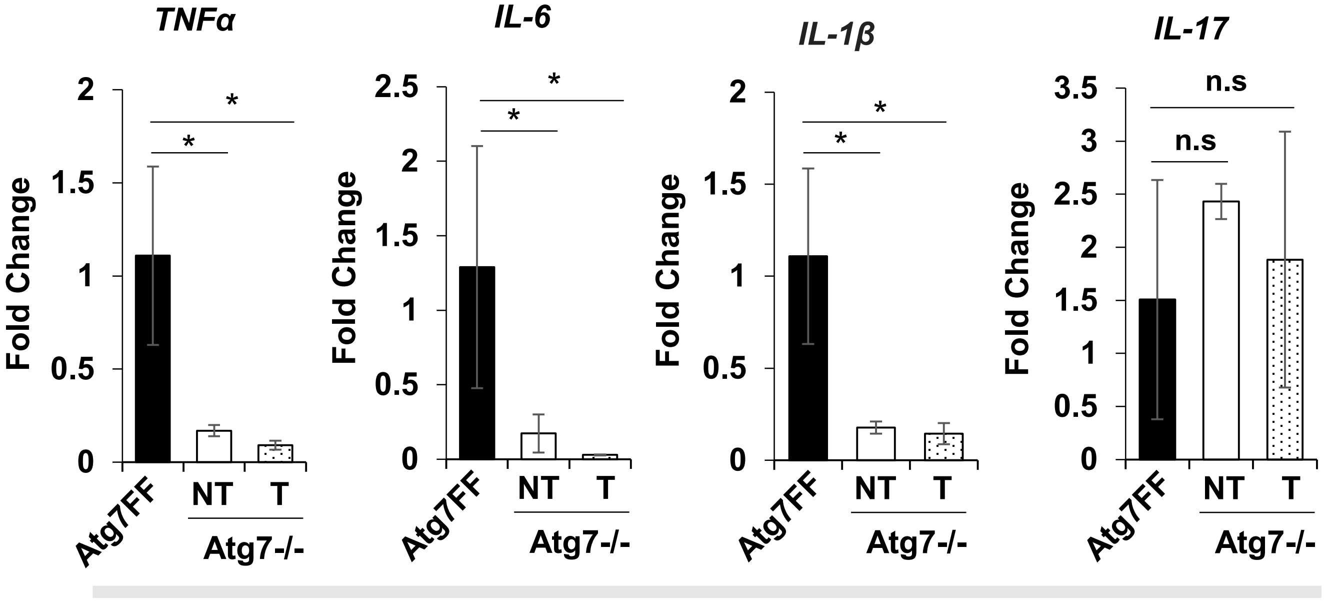


Pro-inflammatory cytokines

### Supplementary Figure S5

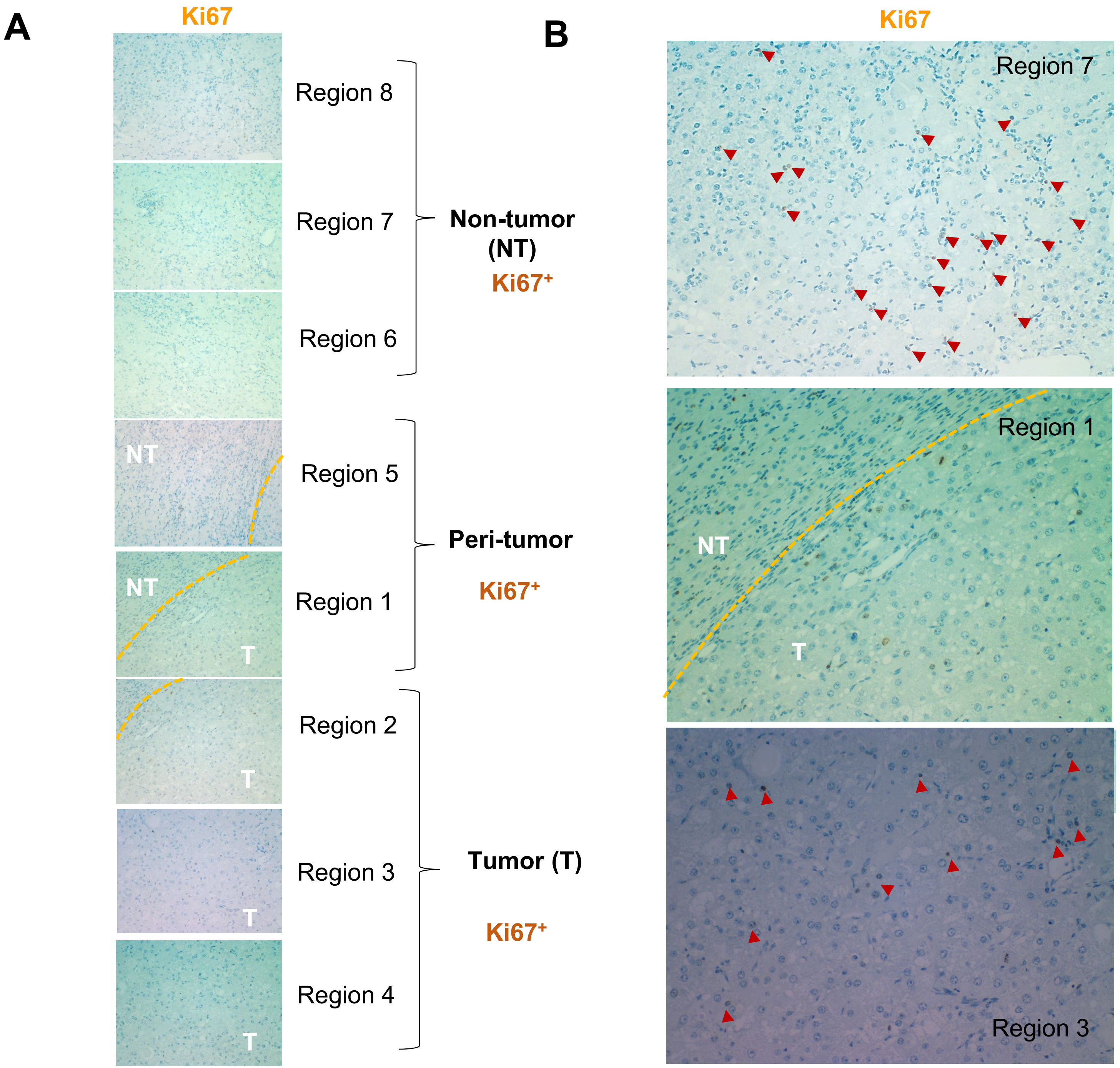

**Supplementary Figure S6**

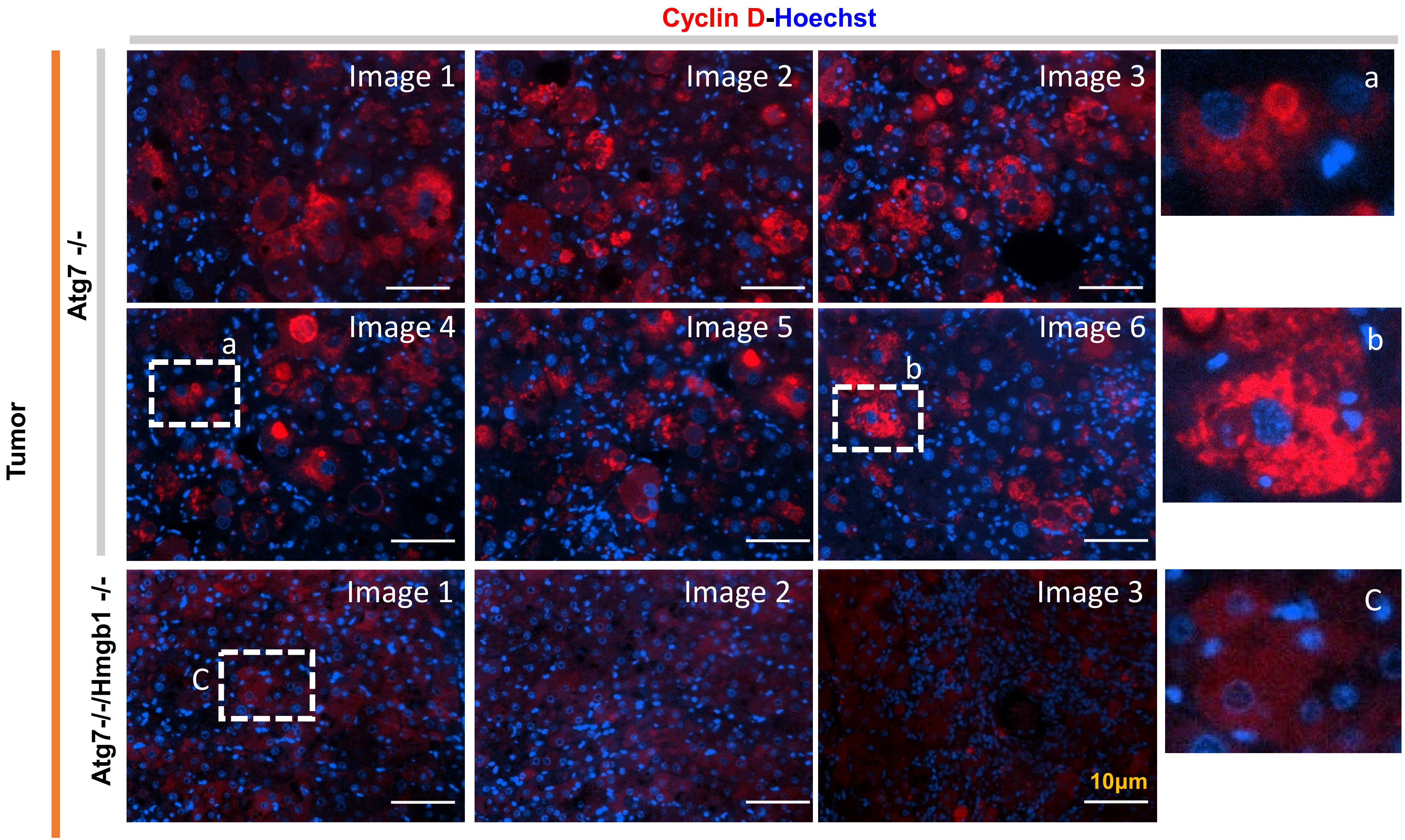

### Supplementary Figure S7

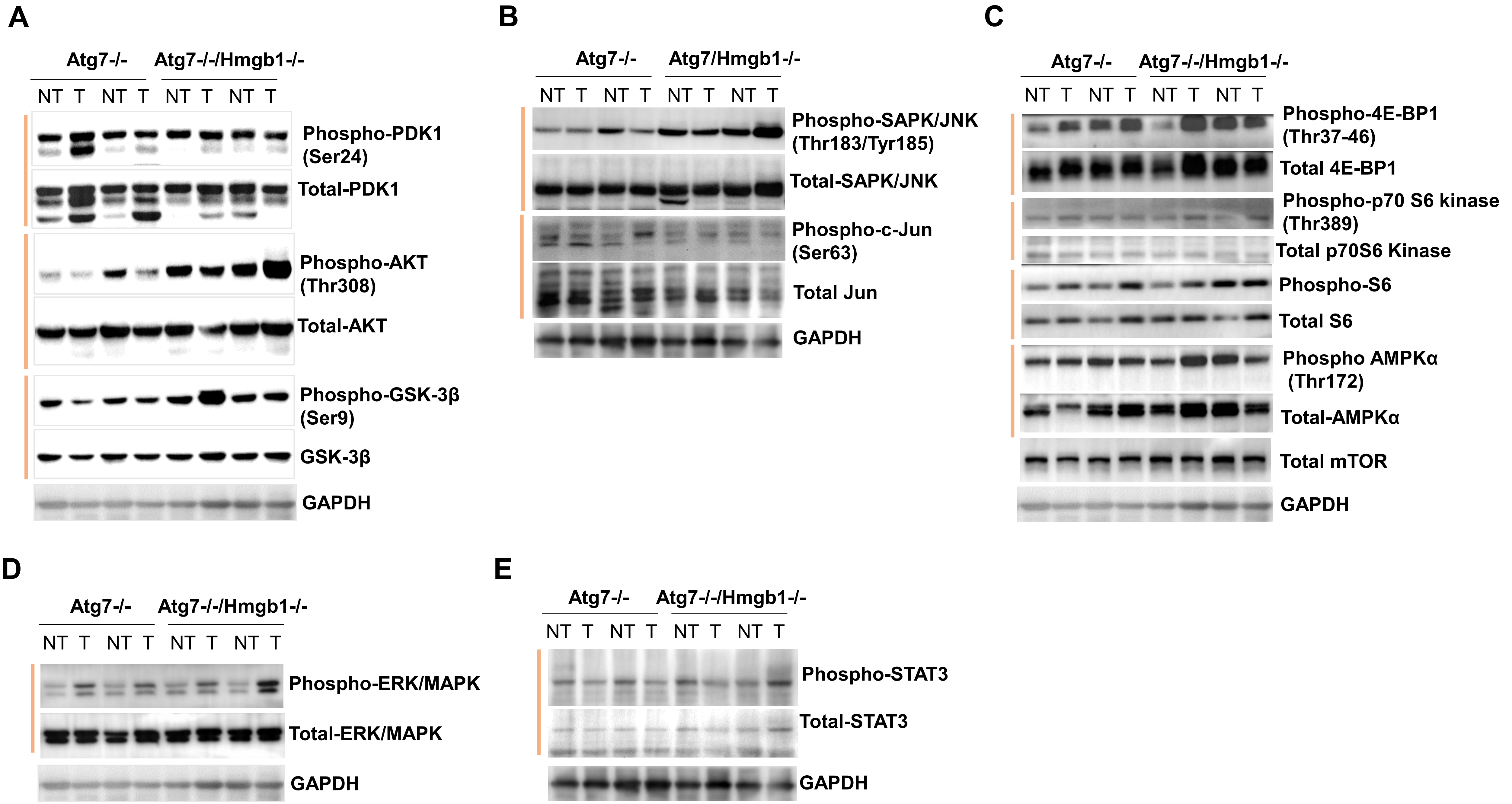

#### Supplementary Figure S8

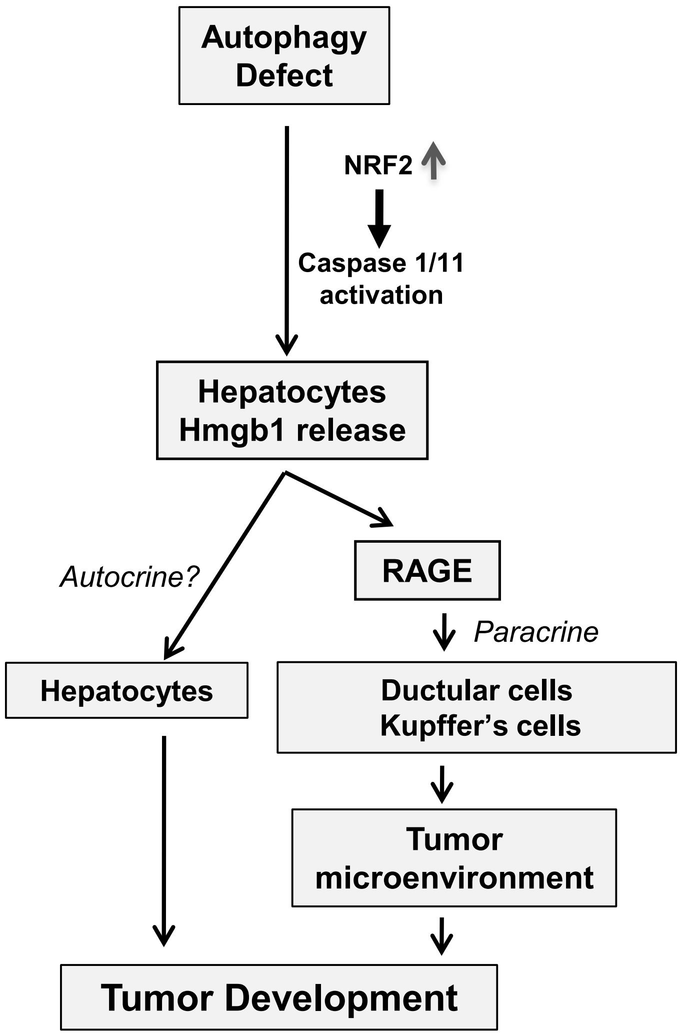

**Supplementary Table S6.**

| Parameter | Non-Tumor(NT) | Peri-tumor | Tumor(T) | Remark |
|---|---|---|---|---|
| SQSTM1 | + | + | + | Autophagy status |
| Ubiquitin(UB) | + | + | + | Autophagy status |
| HNF4α | + | + | + | Hepatocyte marker |
| CK19 | + | + | - | Ductular cell |
| SOX9 | + | + | - | Ductular cell |
| F4/80 | + | + | + | Immune cell-Macrophage |
| MPO | + | + | - | Immune cell-Neutrophil |
| CD3 | - | - | - | Immune cell-T cell |
| CD45R | + | + | - | Immune cell-B cell |
| Desmin | + | + | - | Fibroblast |
