## Supplementary Table S1-Gene Lists-Overlapped for "The HMGB1-RAGE axis modulates the growth of autophagy-deficient hepatic tumors"

**Table S1A: List of overlapped genes that are upregulated in tumors of both *Atg7-/-* and *Atg7-/-/Hmgb1-/-* liver**

| Name | Trxid | logFC.A7KO_TumvsA7KO_NT | logFC.A7HDKO_TumvsA7HDKO_NT |
|---|---|---|---|
| | | p<0.01 | p<0.01 |
| **Psat1** | NM_177420 | 3.029041166 | 2.563235808 |
| **Myo5c** | NM_001081322 | 2.907981904 | 3.801311895 |
| **Psph** | NM_133900 | 1.712390884 | 1.092648969 |
| **Tnfrsf10b** | NM_020275 | 1.664617483 | 1.462540758 |
| **Mfsd6** | NM_178081 | 0.97740622 | 1.546146808 |
| **Dnajc10** | NM_024181 | 1.430535621 | 0.942580926 |
| **Itih5** | NM_172471 | 1.124263494 | 1.500130621 |
| **Abcc4** | NM_001033336 | 1.301240593 | 1.830213073 |
| **Fhdc1** | NM_001033301 | 1.96063355 | 1.879074693 |
| **St3gal2** | NM_009179 | 1.229805694 | 2.135258743 |
| **5330417C22Rik** | NM_001033304 | 1.115301609 | 2.392287376 |
| **Agfg1** | NM_010472 | 0.916963403 | 1.284124708 |
| **Npdc1** | NM_008721 | 1.620085834 | 0.870077813 |
| **Gls** | NM_001113383 | 1.316070409 | 1.614709654 |
| **Asns** | NM_012055 | 3.214071574 | 3.593073769 |
| **Slc3a2** | NM_001161413 | 0.933194133 | 0.769339267 |
| **Ptgs2** | NM_011198 | 2.478687942 | 4.908292457 |
| **Erich4** | NM_001039243 | 3.404181804 | 4.120596436 |
| **Slc35f2** | NM_028060 | 2.341472503 | 3.7669027 |
| **Aldh18a1** | NM_019698 | 1.168219442 | 1.17651422 |
| **Zfand2a** | NM_001159908 | 1.094788263 | 1.112111927 |
| **Phf3** | NM_001081080 | 0.599417657 | 0.844386661 |
| **Arfgap3** | NM_025445 | 0.990845045 | 0.850419729 |
| **Ppp4r1** | NM_001114131 | 0.654585693 | 0.899064197 |
| **Ctr9** | NM_009431 | 0.819807407 | 1.058892475 |
| **AI661453** | NM_145489 | 0.841324947 | 0.910323253 |
| **Golm1** | NM_001035122 | 1.332308642 | 1.279528819 |
| **Psrc1** | NM_019976 | 1.978184807 | 1.943295062 |

**Table S1B: List of overlapped genes that are downregulated in tumors of both *Atg7-/-* and *Atg7-/-/Hmgb1-/-* liver**

| Name | Trxid | logFC.A7KO_TumvsA7KO_NT | logFC.A7HDKO_TumvsA7HDKO_NT |
|---|---|---|---|
| | | p<0.01 | p<0.01 |
| Ppp1r1b | NM_144828 | -2.902033604 | -1.730131727 |
| Cpsf4l | NM_029794 | -2.580176407 | -1.999257305 |
| Socs2 | NM_007706 | -1.284769604 | -1.082663765 |
| Tcf21 | NM_011545 | -1.955217378 | -1.414114392 |
| Upp2 | NM_001289660 | -3.132648942 | -1.470266618 |
| Ramp1 | NM_016894 | -1.423867939 | -1.413534634 |
| Fam180a | NM_173375 | -2.556192143 | -1.24903992 |
| Dpep1 | NM_007876 | -2.453861648 | -1.259371734 |
| Cirbp | NM_007705 | -0.71398568 | -0.824116281 |
| Fam47e | NM_001033478 | -3.504793115 | -2.616401195 |
| Sp5 | NM_022435 | -4.199604299 | -1.9564847 |
