## Supplementary Table S2-Gene Lists- Atg7KO only upregulated for "The HMGB1-RAGE axis modulates the growth of autophagy-deficient hepatic tumors"

**Table S2: List of upregulated genes in tumors of *Atg7-/-* liver**

| Name | Trxid | logFC.A7KO_TumvsA7KO_NT |
|---|---|---|
| 1700016C15Rik | NM_027077 | 4.508710694 |
| Bglap3 | NM_001305449 | 5.705775664 |
| Cpe | NM_013494 | 4.057316738 |
| Memo1 | NM_133771 | 1.058618403 |
| Uxs1 | NM_026430 | 1.24417276 |
| Stom | NM_013515 | 1.203096012 |
| Abcc12 | NM_172912 | 2.089560059 |
| Npc1l1 | NM_207242 | 4.082554273 |
| Hpse2 | NM_001081257 | 2.73728667 |
| Klhl13 | NM_026167 | 1.555802729 |
| Pmaip1 | NM_021451 | 2.212173417 |
| St6galnac2 | NM_009180 | 2.263932532 |
| Ormdl1 | NM_145517 | 1.369878186 |
| Slc16a14 | NM_027921 | 5.089052938 |
| Trpv4 | NM_022017 | 2.02965189 |
| Bend6 | NM_177235 | 1.976632945 |
| Bco1 | NM_021486 | 1.821089663 |
| Slc25a4 | NM_007450 | 2.234481551 |
| Gm9926 | NR_040528 | 4.424314207 |
| 9430016H08Rik | NM_001081181 | 0.880136916 |
| Scamp5 | NM_001301635 | 1.226954953 |
| 2410015M20Rik | NM_153152 | 0.911631095 |
| Clic5 | NM_172621 | 1.617124934 |
| Elfn2 | NM_183141 | 1.885579365 |
| G6pdx | NM_008062 | 2.092594807 |
| Mapt | NM_001285454 | 1.492232774 |
| Mrpl30 | NM_027098 | 0.71562573 |
| Rundc3a | NM_016759 | 1.690679495 |
| Atg12 | NM_026217 | 1.167841088 |
| 1110008P14Rik | NM_198001 | 1.097811064 |
| Synm | NM_207663 | 1.989049545 |
| Amn | NM_033603 | 3.171643275 |
| Setd7 | NM_080793 | 1.113630482 |
| Tfcp2l1 | NM_023755 | 1.211866131 |
| Ccser2 | NM_027045 | 0.716053066 |
| Slc6a8 | NM_001142809 | 1.457922232 |
| Ak7 | NM_030187 | 2.028421321 |
| Ostc | NM_025509 | 0.835166339 |
| Wsb2 | NM_021539 | 0.826269039 |
| Aldh1b1 | NM_028270 | 1.141927315 |
| Ddx39 | NM_197982 | 0.815208344 |
| Rbm3 | NM_001166409 | 0.948175016 |
| Lipn | NM_027340 | 3.391957938 |
| Chka | NM_013490 | 0.883361975 |
| Reep5 | NM_007874 | 1.046478999 |
| Ptgr1 | NM_025968 | 2.024843193 |
| Lsg1 | NM_178069 | 0.651883688 |
| Tex30 | NM_029368 | 1.278872817 |
| Hist1h2bc | NM_023422 | 1.25998837 |
| Fam162a | NM_027342 | 1.03657704 |
| Uqcc2 | NM_026063 | 1.259565535 |
| Skp2 | NM_001285980 | 1.232227197 |
| Bloc1s2 | NM_028607 | 0.871882374 |
| Enpp5 | NM_032003 | 1.046732324 |
| Meig1 | NM_008579 | 2.83224109 |
| Hilpda | NM_001190461 | 1.470611263 |

| | | |
|---|---|---|
| Gla | NM_013463 | 1.14366239 |
| Hist1h4h | NM_153173 | 2.095782897 |
| Comtd1 | NM_026965 | 1.406568889 |
| Smpdl3b | NM_133888 | 2.348407146 |
| Homer3 | NM_001146153 | 2.044305689 |
| Dusp4 | NM_176933 | 1.414944502 |
| Fam81a | NM_029784 | 2.144760769 |
| Fam98a | NM_133747 | 0.843771751 |
| Ces1a | NM_001013764 | 2.719160744 |
| Gnl3 | NM_153547 | 0.86011413 |
| Col22a1 | NM_027174 | 2.681694833 |
| Cyp4a12a | NM_177406 | 3.107042572 |
| Dpf1 | NM_013874 | 1.656049563 |
| Pfkp | NM_001291071 | 2.487601678 |
| Acsm2 | NM_146197 | 3.476103345 |
| Timm8b | NM_013897 | 0.674314765 |
| Atp11b | NM_029570 | 0.644528785 |
| 2310034O05Rik | NR_040679 | 3.030583289 |
| Mrps10 | NM_001146211 | 1.262234103 |
| Cml5 | NM_023493 | 2.861632364 |
| H2-T10 | NR_046286 | 1.262103419 |
| Ppcs | NM_026494 | 0.634904088 |
| Plin3 | NM_025836 | 1.096992207 |
| Neu1 | NM_010893 | 1.54770784 |
| Nmrk1 | NM_145497 | 0.912599212 |
| Zfp706 | NM_026521 | 0.791228956 |
| Kazn | NM_001109685 | 1.844419097 |
| Nif3l1 | NM_022988 | 0.780507533 |
| Cox19 | NM_197980 | 0.940311488 |
| Rnf11 | NM_013876 | 0.689667331 |
| Miox | NM_019977 | 3.906782985 |
| Xkr9 | NM_001011873 | 1.082106033 |
| Eif4e | NM_007917 | 0.616940865 |
| Mea1 | NM_010787 | 1.249724064 |
| Mff | NM_029409 | 0.586504793 |
| Usp50 | NM_029163 | 2.32673094 |
| Imp4 | NM_178601 | 0.589544434 |
| Nme2 | NM_008705 | 0.863973852 |
| Tomm20 | NM_024214 | 0.713920361 |
| Cnih1 | NM_009919 | 0.85380939 |
| Crym | NM_016669 | 1.760306505 |
| Rnf128 | NM_001254761 | 0.87907497 |
| Ssu72 | NM_026899 | 0.651710244 |
| Strbp | NM_009261 | 0.904679094 |
| Ndufa11 | NM_027244 | 0.947078947 |
| Trim35 | NM_029979 | 1.49873895 |
| Chp2 | NM_027363 | 1.256670399 |
| Sdhaf4 | NM_026503 | 0.805005634 |
| Ermp1 | NM_001081213 | 0.724066035 |
| Fam96a | NM_026635 | 0.73115171 |
| Ero1l | NM_015774 | 0.946590504 |
| Tmbim1 | NM_027154 | 0.837770826 |
| Gclc | NM_010295 | 1.181609684 |
| Hrasls | NM_013751 | 5.666431261 |
| Ttc16 | NM_001290563 | 2.105028051 |
| Zdhhc2 | NM_178395 | 1.558344597 |
| Emp2 | NM_007929 | 0.796447365 |
| Tes | NM_207176 | 1.815729931 |
| Abcg2 | NM_011920 | 1.403239608 |

| | | |
|---|---|---|
| Slc16a6 | NM_001029842 | 0.986316164 |
| Lgals1 | NM_008495 | 1.541007168 |
| Txndc9 | NM_172054 | 0.854990129 |
| Cgref1 | NM_026770 | 1.936503323 |
| Chid1 | NM_001142681 | 0.660394177 |
| Eif2s1 | NM_026114 | 0.565318229 |
| Cystm1 | NM_001081365 | 1.825962798 |
| Il17rb | NM_019583 | 0.800959816 |
| Nrg1 | NM_178591 | 1.494835983 |
| Atad2 | NM_027435 | 0.72091328 |
| Coprs | NM_025556 | 1.558198964 |
| Cops5 | NM_001277101 | 0.681672505 |
| Tceb1 | NM_026456 | 0.882079634 |
| E230025N22Rik | NM_172831 | 3.142837365 |
| Creg1 | NM_011804 | 1.047236195 |
| Ugt2b35 | NM_172881 | 1.125915899 |
| Loxl2 | NM_033325 | 1.838328401 |
| Rtn4r | NM_022982 | 2.915770429 |
| Rian | NR_028261 | 5.074759668 |
| Ccne1 | NM_007633 | 1.40924422 |
| Aida | NM_181732 | 1.039220143 |
| Prph | NM_001163589 | 3.555486708 |
| C920025E04Rik | NM_001271005 | 1.705431003 |
| Atg4a | NM_174875 | 0.715496683 |
| Atp6v1g1 | NM_024173 | 0.667579147 |
| Tyw5 | NM_001037742 | 0.973080092 |
| Eif4ebp1 | NM_007918 | 0.919404085 |
| Pgap2 | NM_001291358 | 0.624530057 |
| Pla2g2e | NM_012044 | 6.126929708 |
| Tcea1 | NM_011541 | 0.602157345 |
| Ppap2c | NM_015817 | 0.929603478 |
| Gdf15 | NM_011819 | 1.47471061 |
| Rogdi | NM_133185 | 0.956674691 |
| Cnep1r1 | NM_029074 | 0.59019165 |
| Mlkl | NM_029005 | 0.664083525 |
| Med21 | NM_025315 | 1.021929834 |
| Ranbp1 | NM_011239 | 0.590016095 |
| Gpc1 | NM_016696 | 1.082331941 |
| Morc1 | NM_010816 | 3.967199304 |
| 4933434E20Rik | NM_001287087 | 0.585143739 |
| Rars | NM_025936 | 0.724501143 |
| Neurl1a | NM_021360 | 1.386080889 |
| Ercc3 | NM_133658 | 0.793524972 |
| Fastkd2 | NM_172422 | 0.688846491 |
| Chaf1a | NM_013733 | 1.263388708 |
| Klhdc2 | NM_027117 | 0.68930879 |
| Degs1 | NM_007853 | 0.786915362 |
| Cycs | NM_007808 | 0.624986311 |
| Fbxo25 | NM_025785 | 0.894687425 |
| Ypel5 | NM_027166 | 0.645241638 |
| Pdilt | NM_027943 | 1.031570128 |
| Sbsn | NM_172205 | 1.734756983 |
| Gdpd1 | NM_025638 | 2.098056132 |
| Tmem54 | NM_025452 | 5.430250857 |
| H2-Bl | NM_008199 | 1.747526221 |
| Vdr | NM_009504 | 2.588532053 |
| Trim61 | NM_001177551 | 2.901623502 |
| Syt14 | NM_001301370 | 2.356711989 |
| Cdc34 | NM_177613 | 0.740126504 |

| | | |
|---|---|---|
| Tmem41a | NM_025693 | 0.984906594 |
| Card14 | NM_130886 | 2.401166515 |
| Unc50 | NM_026123 | 0.726796873 |
| Dcun1d5 | NM_029775 | 0.566567141 |
| Arhgap27 | NM_001205236 | 0.760867225 |
| Bid | NM_007544 | 0.820852818 |
| Cyp39a1 | NM_018887 | 1.523224828 |
| Nip7 | NM_001164472 | 0.697153784 |
| Itgb1bp1 | NM_008403 | 0.785971613 |
| Mdm2 | NM_010786 | 0.596691902 |
| Nudt5 | NM_016918 | 0.683752858 |
| Trmt10a | NM_175389 | 0.907549567 |
| Kif26a | NM_001097621 | 1.617350382 |
| Zdhhc16 | NM_023740 | 0.597070091 |
| Slirp | NM_026958 | 0.716296297 |
| N6amt2 | NM_026526 | 0.870959246 |
| Prkaa1 | NM_001013367 | 0.815736166 |
| Mob4 | NM_025283 | 0.701675186 |
| Tmem37 | NM_019432 | 0.783432303 |
| Cox5b | NM_009942 | 0.74765634 |
| Spink1 | NM_009258 | 2.32312065 |
| Mgarp | NM_026358 | 3.670762159 |
| Cyp2f2 | NM_007817 | 1.187366777 |
| Nabp1 | NM_028696 | 1.746499761 |
| Dhrs9 | NM_175512 | 0.878393584 |
| Bak1 | NM_007523 | 0.848776998 |
| Prelid2 | NM_029942 | 1.40870688 |
| A4galt | NM_001004150 | 2.178949585 |
| Fbxo4 | NM_134099 | 0.715221969 |
| Tomm5 | NM_001099675 | 0.627857448 |
| 2900011O08Rik | NM_144518 | 1.855006724 |
| Pgm3 | NM_001163746 | 0.589142675 |
| Mreg | NM_001005423 | 0.730475399 |
| Cryl1 | NM_030004 | 1.26134356 |
| 2310045N01Rik | NR_132132 | 0.508973153 |
| Mrpl13 | NM_026759 | 0.753347983 |
| Chaf1b | NM_028083 | 0.954841587 |
| Sh3gl1 | NM_013664 | 0.547911286 |
| Cep76 | NM_001081073 | 1.089028927 |
| Apom | NM_018816 | 1.428382397 |
| Rps27l | NM_026467 | 0.660801285 |
| Timm23 | NM_016897 | 0.673565574 |
| Hrsp12 | NM_008287 | 1.04992273 |
| Hk2 | NM_013820 | 0.886566195 |
| Rpl7l1 | NM_025433 | 0.936507508 |
| Vil1 | NM_009509 | 4.701739156 |
| Pla2g12b | NM_023530 | 0.831139773 |
| Slc35b2 | NM_028662 | 0.98041743 |
| Rragd | NM_027491 | 1.41677299 |
| Mrps18a | NM_026768 | 1.117080035 |
| Lamtor3 | NM_019920 | 0.763782125 |
| Mpp6 | NM_001164733 | 0.576423733 |
| Bloc1s6 | NM_019788 | 0.618327448 |
| Mydgf | NM_080837 | 0.63410965 |
| 2610203C22Rik | NR_015470 | 1.431187217 |
| Pbdc1 | NM_001281871 | 0.784048801 |
| Cebpe | NM_207131 | 1.578928467 |
| Serpinb1a | NM_025429 | 1.286209296 |
| Samd4 | NM_001163433 | 0.994865587 |

| | | |
|---|---|---|
| Tmem208 | NM_025486 | 0.747716915 |
| Lacc1 | NM_172488 | 0.821312149 |
| Srpk1 | NM_016795 | 0.621097925 |
| Plcl2 | NM_013880 | 0.713560671 |
| Hsd17b13 | NM_001163486 | 0.899774306 |
| Gprc5b | NM_022420 | 1.186177121 |
| Dnajb9 | NM_013760 | 0.644318981 |
| Atf4 | NM_009716 | 0.555356758 |
| Gm3776 | NM_001243092 | 1.405251262 |
| Wfdc3 | NM_027961 | 2.132262144 |
| Gm14137 | NM_001039223 | 2.937729578 |
| Nars | NM_001142950 | 0.680748934 |
| Rpl31 | NM_001252218 | 0.774026511 |
| Anapc16 | NM_025514 | 0.569678355 |
| Zfr | NM_011767 | 0.637271519 |
| Ankrd9 | NM_175207 | 1.03604233 |
| Slc48a1 | NM_026353 | 1.185348115 |
| Tgif1 | NM_001164074 | 0.634750728 |
| Timm17a | NM_011590 | 0.860999306 |
| A930017M01Rik | NR_033609 | 2.75615252 |
| Trpm6 | NM_153417 | 2.252277653 |
| Fos | NM_010234 | 0.942427989 |
| Trnau1ap | NM_027925 | 0.547811258 |
