## Supplementary Table S3-Gene Lists- Atg7HMGB1DKO only upregulated for "The HMGB1-RAGE axis modulates the growth of autophagy-deficient hepatic tumors"

**Table S3: List of upregulated genes in tumors of *Atg7-/-/Hmgb1-/-* liver**

| Name | Trxid | logFC.A7HDKO_TumvsA7HDKO_NT |
|---|---|---|
| Igf2bp3 | NM_023670 | 4.01146651 |
| Prkar2b | NM_011158 | 2.162894044 |
| Sgsm1 | NM_001254731 | 2.337219859 |
| Tspan8 | NM_146010 | 2.402331505 |
| Eps8l3 | NM_133867 | 1.949271702 |
| Smurf1 | NM_029438 | 1.228415691 |
| Thbs1 | NM_011580 | 1.525201309 |
| Myof | NM_001099634 | 1.98117925 |
| Ppp1r42 | NR_110971 | 3.225887704 |
| Fosl1 | NM_010235 | 2.789566169 |
| Col4a2 | NM_009932 | 1.777820447 |
| Mki67 | NM_001081117 | 1.1394145 |
| Hmgcll1 | NM_173731 | 4.618900452 |
| Trrap | NM_001081362 | 1.477646041 |
| Piezo1 | NM_001037298 | 1.259558832 |
| Loxl4 | NM_053083 | 2.260012986 |
| Aldh1l2 | NM_153543 | 3.251771986 |
| Peak1 | NM_172924 | 1.471759586 |
| Epb4.1l1 | NM_013510 | 1.62888783 |
| Cgnl1 | NM_026599 | 1.34345189 |
| Phgdh | NM_016966 | 2.034169153 |
| Creb3l2 | NM_178661 | 1.359223613 |
| Tmem150c | NM_182841 | 2.183741554 |
| Lnx2 | NM_080795 | 1.140633436 |
| Akap9 | NM_194462 | 1.25092998 |
| Clip2 | NM_009990 | 1.375607643 |
| Plec | NM_011117 | 1.263490438 |
| Kif5c | NM_008449 | 2.483371975 |
| Tspan18 | NM_183180 | 1.784945572 |
| Zmat3 | NM_009517 | 1.381303679 |
| Gtf2ird1 | NM_001081463 | 1.177471155 |
| Mst1r | NM_009074 | 2.719680661 |
| Dusp5 | NM_001085390 | 1.923922419 |
| Fbn1 | NM_007993 | 1.129117446 |
| Col4a1 | NM_009931 | 1.309570157 |
| Cdhr2 | NM_001033364 | 4.217856083 |
| Vill | NM_001164567 | 1.313252814 |
| Tjp2 | NM_011597 | 1.267768071 |
| Ptgfrn | NM_011197 | 1.347288393 |
| Lamc1 | NM_010683 | 1.722005573 |
| Atp8a2 | NM_015803 | 3.505039972 |
| Mical2 | NM_001193305 | 1.260108748 |
| Smad3 | NM_016769 | 1.544048795 |
| Slc39a4 | NM_028064 | 1.704606827 |
| Inhbb | NM_008381 | 2.37787459 |
| Src | NM_009271 | 1.251591704 |
| Fam129b | NM_146119 | 1.092159674 |
| Tgfa | NM_031199 | 1.34581215 |

| | | |
|---|---|---|
| Capn5 | NM_007602 | 1.108907361 |
| Ptpn14 | NM_008976 | 2.229020043 |
| Ppl | NM_008909 | 1.283043782 |
| Vgll3 | NM_028572 | 3.845185063 |
| Abcb1a | NM_011076 | 1.57805285 |
| Nid1 | NM_010917 | 1.680653741 |
| Ankib1 | NM_001289527 | 1.100760337 |
| Steap1 | NM_027399 | 2.358779287 |
| Clcf1 | NM_001310039 | 1.275663678 |
| Nol4l | NM_001134300 | 1.700984775 |
| Fndc3b | NM_173182 | 1.411447463 |
| Zfp57 | NM_001013745 | 2.744548412 |
| Prrc2c | NM_001081290 | 1.177693734 |
| Cep192 | NM_027556 | 1.266741586 |
| Cd93 | NM_010740 | 1.583405559 |
| Cdk6 | NM_009873 | 1.218153092 |
| Abl2 | NM_001136104 | 1.228906997 |
| Kmt2a | NM_001081049 | 1.341456395 |
| Gcn1l1 | NM_172719 | 1.309153652 |
| Mast4 | NM_175171 | 1.450474014 |
| Ppfibp1 | NM_026221 | 1.137027021 |
| Atp2b4 | NM_001167949 | 1.767029283 |
| Ptpn11 | NM_001109992 | 0.952580175 |
| Ehf | NM_007914 | 2.026746783 |
| Slco2a1 | NM_033314 | 1.766033397 |
| Ncor2 | NM_001253905 | 1.223644351 |
| Sema3e | NM_011348 | 4.937159569 |
| Lama5 | NM_001081171 | 2.085026729 |
| Ahnak | NM_001039959 | 1.895332253 |
| Ago2 | NM_153178 | 1.722513187 |
| Mcam | NM_023061 | 1.026002527 |
| Clip1 | NM_001291229 | 0.958073504 |
| Ctps | NM_016748 | 1.168206313 |
| Birc6 | NM_007566 | 1.081240155 |
| Tet3 | NM_183138 | 1.220349997 |
| Fermt1 | NM_198029 | 1.149957212 |
| Myh9 | NM_022410 | 1.244411475 |
| Asap2 | NM_001135192 | 1.337859563 |
| Dhx37 | NM_203319 | 1.118336418 |
| Wscd1 | NM_177618 | 2.284064762 |
| Tead1 | NM_009346 | 1.528066959 |
| Spen | NM_019763 | 1.198344972 |
| Afp | NM_007423 | 5.6170979 |
| Fat1 | NM_001081286 | 1.530507727 |
| Abcb1b | NM_011075 | 1.638734301 |
| Lrrc55 | NM_001033346 | 1.999026756 |
| Spire2 | NM_172287 | 1.57343972 |
| Xpo5 | NM_028198 | 0.972716057 |
| Col4a3 | NM_007734 | 2.646942071 |
| 1700017B0 | NM_028820 | 0.949519007 |

| | | |
|---|---|---|
| Cdc25a | NM_007658 | 1.490297102 |
| Osbpl3 | NM_001163645 | 1.005972593 |
| Emcn | NM_001163522 | 1.195185237 |
| Ywhag | NM_018871 | 0.866073461 |
| Glg1 | NM_009149 | 1.011318209 |
| Pdlim7 | NM_026131 | 1.075888458 |
| Pak6 | NM_001145854 | 1.550329279 |
| Spire1 | NM_194355 | 1.285776928 |
| Flnb | NM_134080 | 1.109886229 |
| Plekhb2 | NM_145516 | 0.842603839 |
| Myo5a | NM_010864 | 2.052346409 |
| Gm15800 | NM_181421 | 1.128937979 |
| Lars | NM_134137 | 0.970690497 |
| Abcc1 | NM_008576 | 1.458025709 |
| Ank | NM_020332 | 0.969561041 |
| Zfp704 | NM_133218 | 1.317260635 |
| Atp6v0a4 | NM_080467 | 1.78893584 |
| Vash1 | NM_177354 | 1.415719956 |
| Zkscan1 | NM_133906 | 0.888311202 |
| Cd276 | NM_133983 | 1.444592747 |
| Lmtk2 | NM_001081109 | 1.324235487 |
| Svil | NM_153153 | 1.006236745 |
| Nin | NM_001286080 | 1.13479203 |
| Ltbp3 | NM_008520 | 1.620639967 |
| Lpp | NM_001145952 | 0.912113252 |
| Ino80d | NM_001081436 | 1.122006324 |
| Trio | NM_001081302 | 1.154338215 |
| Pacs1 | NM_153129 | 1.061142048 |
| Abcc5 | NM_176839 | 2.061812609 |
| Cad | NM_001289522 | 1.244504863 |
| Cep350 | NM_001039184 | 1.068059615 |
| Gldc | NM_138595 | 1.209383684 |
| Mthfd2 | NM_008638 | 1.592621991 |
| Myadm | NM_001093765 | 1.264136897 |
| Rab3d | NM_031874 | 1.059375299 |
| Bhlha15 | NM_010800 | 3.643138169 |
| Col5a2 | NM_007737 | 1.249982113 |
| Rcan3 | NM_022980 | 1.061522845 |
| Fstl3 | NM_031380 | 1.463686742 |
| Arhgap11a | NM_181416 | 1.239274751 |
| Foxk1 | NM_199068 | 1.055445913 |
| Dnm3 | NM_001038619 | 3.448374098 |
| Sbno1 | NM_001081203 | 0.762383692 |
| Clasp1 | NM_001293301 | 1.195660101 |
| Tanc2 | NM_181071 | 1.55729161 |
| Sez6l2 | NM_144926 | 1.914073999 |
| Ldlrad3 | NM_178886 | 1.374312786 |
| Tpr | NM_133780 | 0.90090803 |
| Col15a1 | NM_009928 | 1.175430816 |
| Dip2a | NM_001081419 | 1.400273226 |

| | | |
|---|---|---|
| Numbl | NM_010950 | 1.925340989 |
| Smarcc1 | NM_009211 | 0.86105146 |
| Pycr1 | NM_144795 | 1.754319813 |
| Ap1s3 | NM_183027 | 1.454652884 |
| Dst | NM_133833 | 1.336365799 |
| Plxna2 | NM_008882 | 1.486443897 |
| Kmt2d | NM_001033276 | 1.123211608 |
| Asap1 | NM_010026 | 0.83784541 |
| Gsto1 | NM_010362 | 1.416778529 |
| Aplnr | NM_011784 | 1.803611048 |
| Plxna1 | NM_008881 | 1.139668332 |
| Herc2 | NM_010418 | 1.247801641 |
| Agrn | NM_021604 | 0.965324611 |
| Fam208b | NM_134063 | 1.523460009 |
| Olr1 | NM_138648 | 2.281582057 |
| Nrxn2 | NM_001205234 | 1.456733568 |
| Sort1 | NM_001271599 | 1.094707395 |
| Fnbp1l | NM_001114665 | 1.25473053 |
| Myzap | NM_001033208 | 1.543131622 |
| Kctd3 | NM_172650 | 1.141337963 |
| Nes | NM_016701 | 1.366121593 |
| Map4 | NM_001205332 | 0.968211897 |
| Syne1 | NM_153399 | 1.187130676 |
| Tanc1 | NM_198294 | 1.102297872 |
| Hspg2 | NM_008305 | 0.983955288 |
| Vcl | NM_009502 | 1.066634055 |
| Cachd1 | NM_198037 | 1.826881751 |
| Dhx9 | NM_007842 | 0.793039619 |
| Setx | NM_198033 | 1.174444304 |
| Ubqln2 | NM_018798 | 1.216781593 |
| Tulp4 | NM_054040 | 1.315288361 |
| Fbxo21 | NM_145564 | 1.247478774 |
| Msantd3 | NM_001145924 | 1.608766761 |
| Atr | NM_019864 | 1.065037336 |
| Vars | NM_011690 | 0.665873173 |
| Ly75 | NM_013825 | 2.178737102 |
| Prrg3 | NM_001303028 | 1.391315124 |
| Rapgef6 | NM_175258 | 0.940258067 |
| Mdn1 | NM_001081392 | 1.187055231 |
| Cpd | NM_007754 | 0.800363949 |
| Nipbl | NM_201232 | 0.812890966 |
| Tnrc6b | NM_177124 | 0.840041711 |
| Rai14 | NM_001166408 | 0.804705825 |
| Herc3 | NM_028705 | 1.270888591 |
| Myh14 | NM_001271540 | 1.54097731 |
| Lpl | NM_008509 | 1.681189409 |
| Tnfaip2 | NM_009396 | 0.922386407 |
| Zfp697 | NM_172863 | 2.064057371 |
| Cd200 | NM_010818 | 1.198780108 |
| Nxn | NM_008750 | 1.366094883 |

| | | |
|---|---|---|
| Slc44a3 | NM_145394 | 1.136407326 |
| Notch1 | NM_008714 | 0.918329093 |
| Usp20 | NM_028846 | 0.90509762 |
| BC030870 | NR_033217 | 3.770407512 |
| Ces2c | NM_145603 | 1.245890131 |
| Tcerg1 | NM_001289526 | 0.871098584 |
| Tmem248 | NM_027854 | 0.758441423 |
| Cep170 | NM_001099637 | 1.217622156 |
| Eif3b | NM_133916 | 0.877598627 |
| Dync1h1 | NM_030238 | 1.039223286 |
| Ttc21b | NM_001290669 | 1.291687394 |
| Stc1 | NM_009285 | 2.149851533 |
| Trim56 | NM_201373 | 1.315958926 |
| Zc3hav1l | NM_172467 | 2.014269016 |
| Ednrb | NM_001136061 | 0.99939783 |
| Cdr2 | NM_007672 | 2.101839094 |
| Atxn2 | NM_009125 | 0.930609135 |
| Bicd2 | NM_029791 | 0.940050886 |
| Micall1 | NM_177461 | 0.766545402 |
| Golga4 | NM_018748 | 1.295851389 |
| Prrc2b | NM_001159634 | 1.160124603 |
| Utrn | NM_011682 | 1.262419573 |
| Tjp1 | NM_001163574 | 1.20856165 |
| Rictor | NM_030168 | 1.25129603 |
| Dusp18 | NM_173745 | 1.823449199 |
| Sptan1 | NM_001177667 | 1.240644882 |
| Cttn | NM_001252572 | 0.878504826 |
| Utp20 | NM_175158 | 0.989970357 |
| Rin1 | NM_145495 | 1.753688213 |
| Ckap5 | NM_029437 | 0.874665098 |
| Psd3 | NM_177698 | 0.973135797 |
| Atp2a2 | NM_001110140 | 0.853446486 |
| Wnk1 | NM_198703 | 0.983165902 |
| Setd8 | NM_030241 | 0.726116854 |
| Cers6 | NM_172856 | 1.454599086 |
| Gfpt1 | NM_013528 | 1.557634969 |
| Map4k4 | NM_001252202 | 0.895077303 |
| Zfp106 | NM_011743 | 1.005282048 |
| Tgfbr2 | NM_009371 | 0.893146799 |
| Vps13c | NM_177184 | 1.093660913 |
| Aff1 | NM_133919 | 1.017525269 |
| Ngf | NM_013609 | 1.405931808 |
| B4galt6 | NM_019737 | 1.107827118 |
| Mboat1 | NM_153546 | 2.419017148 |
| Trip12 | NM_133975 | 0.878226822 |
| Rora | NM_001289916 | 0.919776999 |
| 9330102E0 | NR_077223 | 1.966246847 |
| Tubb2b | NM_023716 | 1.67876456 |
| Ttc37 | NM_001081352 | 0.857997073 |
| Abca7 | NM_013850 | 0.792754583 |

| | | |
|---|---|---|
| Xrn1 | NM_011916 | 1.288063535 |
| Mvp | NM_080638 | 0.75762897 |
| Spg11 | NM_145531 | 0.802708765 |
| Tnks | NM_175091 | 1.25761164 |
| Col4a4 | NM_007735 | 2.379203796 |
| Tmem263 | NM_001013028 | 1.142621601 |
| Ern1 | NM_023913 | 1.303246913 |
| Tie1 | NM_011587 | 0.949893374 |
| Aak1 | NM_001040106 | 1.08873326 |
| Itpr3 | NM_080553 | 1.453761897 |
| R3hdm1 | NM_181750 | 0.763584077 |
| Pja2 | NM_001025309 | 0.956089268 |
| Rab11fip5 | NM_177466 | 1.390270536 |
| Arrb1 | NM_178220 | 1.034451189 |
| Zbed6 | NM_001166552 | 1.712607136 |
| Disp1 | NM_001278218 | 1.347516007 |
| Dip2b | NM_172819 | 0.820642783 |
| Ubr4 | NM_001160319 | 1.021165231 |
| Ick | NM_019987 | 0.919415397 |
| Cdh1 | NM_009864 | 1.463093268 |
| Snrnp200 | NM_177214 | 1.027809485 |
| Tor4a | NM_146115 | 0.913831025 |
| Hivep1 | NM_007772 | 1.193976793 |
| Sptbn1 | NM_175836 | 1.035578882 |
| Ets2 | NM_011809 | 0.801959659 |
| Mllt4 | NM_010806 | 0.828193212 |
| Heg1 | NM_175256 | 1.023138344 |
| Baz2a | NM_054078 | 0.901044531 |
| Nid2 | NM_008695 | 1.848809938 |
| Smchd1 | NM_028887 | 0.770074174 |
| Cdkn2b | NM_007670 | 1.338994763 |
| Notch4 | NM_010929 | 1.136935384 |
| Hectd2 | NM_172637 | 1.386227196 |
| Slc7a1 | NM_007513 | 0.991439968 |
| Prkca | NM_011101 | 1.120211356 |
| Cul9 | NM_001081335 | 0.95207628 |
| Plxnd1 | NM_026376 | 0.764952568 |
| Igf2 | NM_001122736 | 7.682743649 |
| Irs1 | NM_010570 | 1.156802254 |
| Kntc1 | NM_001042421 | 1.402396936 |
| Cdkn1a | NM_001111099 | 0.722202749 |
| Morc4 | NM_029413 | 1.690057626 |
| Ggt1 | NM_001305992 | 3.165646461 |
| Srrm2 | NM_175229 | 0.897663037 |
| Ehd4 | NM_133838 | 0.755519585 |
| Shtn1 | NM_175172 | 0.732489646 |
| Gltscr1l | NM_001100452 | 1.055709308 |
| Als2cl | NM_001146060 | 0.892894703 |
| Tnfrsf11a | NM_009399 | 1.054697093 |
| Bcl2l11 | NM_009754 | 1.05183982 |

| | | |
|---|---|---|
| Nktr | NM_010918 | 0.8670041 |
| Ubn2 | NM_177185 | 0.886410612 |
| Edem3 | NM_001039644 | 0.88522699 |
| Ep300 | NM_177821 | 1.149717546 |
| Kif3c | NM_008445 | 1.365107383 |
| Mid2 | NM_011845 | 1.568783081 |
| Pdgfb | NM_011057 | 1.138461538 |
| Ppip5k2 | NM_173760 | 1.12025512 |
| Garem | NM_001033445 | 0.811428388 |
| Sh3pxd2a | NM_001164717 | 0.998346845 |
| Ncoa6 | NM_019825 | 0.98417897 |
| Pacsin2 | NM_001159509 | 1.010864206 |
| Tpp2 | NM_009418 | 0.945446909 |
| Slc2a1 | NM_011400 | 0.706897452 |
| Slc25a43 | NM_001085497 | 1.537781834 |
| Kif2a | NM_001145779 | 0.803637411 |
| Ash1l | NM_138679 | 1.148964983 |
| Sash1 | NM_175155 | 0.806789986 |
| Dmxl1 | NM_001081371 | 1.142353451 |
| Hipk1 | NM_001301304 | 0.967124496 |
| Lyst | NM_010748 | 1.060988363 |
| Tacc2 | NM_206856 | 1.110300529 |
| Diap2 | NM_172493 | 1.003620446 |
| Mturn | NM_001289741 | 1.971273673 |
| Apaf1 | NM_001282947 | 0.895135101 |
| Arfgef2 | NM_001085495 | 0.74742076 |
| Elk4 | NM_007923 | 0.913389782 |
| Fbxw8 | NM_172721 | 0.710444679 |
| Igfbp3 | NM_008343 | 1.406194411 |
| Eps8l2 | NM_133191 | 1.246379707 |
| Polr1a | NM_009088 | 0.742536452 |
| Ankhd1 | NM_175375 | 0.749060197 |
| Ubr5 | NM_001112721 | 0.751002406 |
| Abca2 | NM_007379 | 0.754560959 |
| Atp8b1 | NM_001001488 | 0.829496737 |
| Med13l | NM_172424 | 0.985260502 |
| Nedd9 | NM_001111324 | 1.078849562 |
| Jag1 | NM_013822 | 1.291366081 |
| Prr13 | NM_025385 | 0.720514066 |
| Ankrd1 | NM_013468 | 2.267425855 |
| Mycbp2 | NM_207215 | 0.860020012 |
| Sh3pxd2b | NM_177364 | 1.124766602 |
| Arid1b | NM_001085355 | 0.794902486 |
| Smpd3 | NM_021491 | 0.671052276 |
| Wdr26 | NM_145514 | 0.782220181 |
| Cbl | NM_007619 | 1.23121128 |
