## Supplementary Table S4-Gene Lists- Atg7KO only downregulated for "The HMGB1-RAGE axis modulates the growth of autophagy-deficient hepatic tumors"

**Table S4: List of downregulated genes in tumors of *Atg7-/-* liver**

| Name | Trxid | logFC.A7KO_TumvsA7KO_NT |
|---|---|---|
| Fgfr3 | NM_001163215 | -1.657459286 |
| Cntfr | NM_001146080 | -3.515380355 |
| Ntn1 | NM_008744 | -1.992666704 |
| Tmc4 | NM_181820 | -3.306708874 |
| Col27a1 | NM_025685 | -2.36899913 |
| Tbx3 | NM_198052 | -3.053562957 |
| Tshz2 | NM_080455 | -1.671448409 |
| Tbx20 | NM_001205085 | -1.822004108 |
| Tnxb | NM_031176 | -1.964234739 |
| Amotl2 | NM_019764 | -1.053191798 |
| Axin2 | NM_015732 | -2.12645871 |
| Efemp1 | NM_146015 | -1.663747977 |
| Angpt1 | NM_001286062 | -2.317400833 |
| Abcc9 | NM_011511 | -0.865631107 |
| Adam33 | NM_033615 | -3.240806766 |
| Rbms3 | NM_001172121 | -2.003340634 |
| Setbp1 | NM_053099 | -2.569501974 |
| Lgr5 | NM_010195 | -3.331492215 |
| Smim1 | NM_001163721 | -1.823808027 |
| Pdgfra | NM_011058 | -2.143903621 |
| Vamp2 | NM_009497 | -1.157403056 |
| Syt7 | NM_018801 | -2.831655611 |
| Hpse | NM_152803 | -1.274264052 |
| Mamdc2 | NM_174857 | -3.27389078 |
| Lect2 | NM_010702 | -2.010445876 |
| Adgrl1 | NM_181039 | -1.541129031 |
| Pbx1 | NM_001291509 | -2.096594576 |
| Kcnk1 | NM_008430 | -3.572310008 |
| Gabra3 | NM_008067 | -1.660124567 |
| Arhgef17 | NM_001081116 | -1.751581045 |
| Crispld2 | NM_030209 | -2.421581748 |
| Znrf3 | NM_001080924 | -1.044590357 |
| Itga1 | NM_001033228 | -0.919418887 |
| Pcnt | NM_001282992 | -0.963213943 |
| Cyp2c67 | NM_001024719 | -2.299652958 |
| Macc1 | NM_001163136 | -4.48107464 |
| Abi3bp | NM_001014399 | -2.533379653 |
| Il17re | NM_145826 | -2.606362787 |
| Daam2 | NM_001008231 | -2.150765739 |
| Map2 | NM_001039934 | -2.479673424 |
| Fam171b | NM_175514 | -1.598335916 |
| Dennd2a | NM_172477 | -1.427599106 |
| Ltbp4 | NM_175641 | -1.393053496 |
| Smad6 | NM_008542 | -1.690478546 |
| P3h2 | NM_173379 | -2.012157823 |

| | | |
|---|---|---|
| Rai2 | NM_001103367 | -2.385243218 |
| Cyp2e1 | NM_021282 | -4.40516097 |
| Cited4 | NM_019563 | -3.41657477 |
| Rspo3 | NM_028351 | -1.897466737 |
| Prom2 | NM_178047 | -3.042089307 |
| Mtcl1 | NM_172963 | -2.937444205 |
| Slc43a3 | NM_021398 | -1.094342054 |
| Gna14 | NM_008137 | -3.1021797 |
| Ddr2 | NM_022563 | -1.262428542 |
| Hand2 | NM_010402 | -1.831249297 |
| Mmp7 | NM_010810 | -3.992127952 |
| Pon1 | NM_011134 | -2.739483235 |
| Nynrin | NM_001040072 | -1.900284986 |
| Ildr1 | NM_001285788 | -2.173549821 |
| Hnrnph3 | NM_001079824 | -0.963968275 |
| Dlc1 | NM_001194940 | -0.704396605 |
| Srgap3 | NM_080448 | -2.452396022 |
| Plekha6 | NM_001160268 | -1.017830143 |
| C630043F0 | NR_027923 | -2.019867586 |
| Ltbp1 | NM_206958 | -1.991288582 |
| Kcnb1 | NM_008420 | -2.447306091 |
| Dact3 | NM_001081655 | -2.004152457 |
| Cyp26b1 | NM_001177713 | -2.911330822 |
| Adcy1 | NM_009622 | -2.411221129 |
| Tgfbi | NM_009369 | -1.073696496 |
| Avpr1a | NM_016847 | -2.744459415 |
| Il6st | NM_010560 | -1.003159061 |
| Zfp629 | NM_177226 | -1.028608915 |
| Sfrp5 | NM_018780 | -3.664679373 |
| Plekhs1 | NM_001164263 | -3.045503102 |
| Grik3 | NM_001081097 | -4.640588139 |
| Syt9 | NM_021889 | -1.805935912 |
| Slco1b2 | NM_020495 | -1.700821071 |
| Gm16551 | NR_045284 | -2.530913827 |
| Gdf10 | NM_145741 | -2.707061389 |
| Slc22a27 | NM_134256 | -4.334895682 |
| Rab27b | NM_001082553 | -2.309801678 |
| Slc12a8 | NM_001083902 | -2.345049057 |
| Sema3c | NM_013657 | -2.454609545 |
| Dtna | NM_001285807 | -2.875785151 |
| Grhl2 | NM_026496 | -3.415827005 |
| Lrat | NM_023624 | -1.399610447 |
| Per1 | NM_011065 | -1.01607839 |
| Itga8 | NM_001001309 | -1.915329857 |
| AW549542 | NR_045702 | -4.336389365 |
| Ihh | NM_010544 | -3.125886048 |
| Igsf10 | NM_001162884 | -1.976486761 |

| | | |
|---|---|---|
| Lrrc32 | NM_001113379 | -0.833100278 |
| Rgn | NM_009060 | -3.602081026 |
| Pcdhga5 | NM_033588 | -2.322133321 |
| Adssl1 | NM_007421 | -1.261752688 |
| Dcn | NM_001190451 | -1.326653418 |
| Slc1a2 | NM_001077514 | -4.373404003 |
| Robo2 | NM_175549 | -1.531867693 |
| Thbs2 | NM_011581 | -1.613598557 |
| Sema6c | NM_011351 | -2.14820077 |
| Sftpd | NM_009160 | -2.052917252 |
| Smim6 | NM_001162998 | -2.317980635 |
| Wt1 | NM_144783 | -2.998262248 |
| Tns3 | NM_001083587 | -0.684719773 |
| Gcn1l1 | NM_172719 | -0.621353912 |
| Gprasp1 | NM_026081 | -1.118858706 |
| Podn | NM_001285958 | -2.177381744 |
| Csrp3 | NM_001198841 | -2.40104238 |
| Lrrk1 | NM_146191 | -0.921248179 |
| Fzd1 | NM_021457 | -1.257262236 |
| Map3k9 | NM_177395 | -2.210129348 |
| Sprn | NM_183147 | -1.895419603 |
| Hs3st1 | NM_010474 | -1.995476089 |
| Lipg | NM_010720 | -2.224779857 |
| Muc1 | NM_013605 | -2.962488735 |
| Ston1 | NM_029858 | -0.880287279 |
| Olfr1034 | NM_001011872 | -2.490825591 |
| Slc22a17 | NM_021551 | -2.192932934 |
| Shisa3 | NM_001033415 | -3.495535757 |
| Igf1r | NM_010513 | -1.867965831 |
| Pla2r1 | NM_008867 | -1.316868825 |
| Sdk2 | NM_172800 | -1.963831053 |
| Adamtsl2 | NM_029981 | -2.077632187 |
| Bmp5 | NM_007555 | -1.379075141 |
| Capn6 | NM_007603 | -2.67631161 |
| Ikzf4 | NM_011772 | -3.556207953 |
| Zkscan17 | NM_001291014 | -0.85733419 |
| Mycn | NM_008709 | -2.33082479 |
| Ggt5 | NM_011820 | -1.367052866 |
| Efs | NM_010112 | -2.641098475 |
| Fam174b | NM_001162532 | -1.291187057 |
| Slc5a1 | NM_019810 | -2.484653749 |
| Cygb | NM_030206 | -1.596707892 |
| Add3 | NM_001164099 | -0.926094501 |
| Adamtsl1 | NM_029967 | -1.403924774 |
| Gm4980 | NM_001195529 | -1.367643732 |
| Nkd1 | NM_027280 | -1.599029078 |
| Fgf12 | NM_010199 | -2.033123195 |

| | | |
|---|---|---|
| Phf2 | NM_011078 | -0.63690929 |
| Prdm16 | NM_001177995 | -1.499742768 |
| Cyp2b9 | NM_010000 | -4.545015118 |
| Gucy1a3 | NM_021896 | -1.15661142 |
| Nr1d1 | NM_145434 | -1.629363334 |
| Thsd4 | NM_001040426 | -2.631763503 |
| Hivep1 | NM_007772 | -0.749780332 |
| Notum | NM_175263 | -1.277162014 |
| Agtr1a | NM_177322 | -1.3176038 |
| Sox12 | NM_011438 | -1.518030768 |
| Fasn | NM_007988 | -1.988445789 |
| Alox12e | NM_145684 | -4.270887381 |
| Ccdc166 | NM_001163518 | -1.844638548 |
| Slc4a3 | NM_009208 | -2.436241109 |
| Cbfa2t3 | NM_009824 | -1.215599055 |
| Fam46b | NM_175307 | -2.720875566 |
| Nr2f2 | NM_183261 | -0.973216117 |
| Slc22a26 | NM_146232 | -5.068043636 |
| Pam | NM_013626 | -0.908664972 |
| Dzank1 | NM_172859 | -2.058079253 |
| Pygm | NM_011224 | -1.934488939 |
| Emilin1 | NM_133918 | -1.17486056 |
| Ptch1 | NM_008957 | -0.959266061 |
| Nfia | NM_001122953 | -0.718943298 |
| Pclo | NM_011995 | -4.231723716 |
| C7 | NM_001243837 | -3.473406132 |
| Selp | NM_011347 | -1.742414568 |
| Bspry | NM_138653 | -2.455484693 |
| Tnrc18 | NM_178242 | -0.659498925 |
| D630039A( | NM_178727 | -1.496732097 |
| Clca3a1 | NM_009899 | -0.768249755 |
| Plxna4 | NM_175750 | -1.397610051 |
| Scube1 | NM_001271473 | -1.479009813 |
| Fat4 | NM_183221 | -1.418226579 |
| Slc9a7 | NM_177353 | -1.754168283 |
| Col14a1 | NM_181277 | -1.048891461 |
| Ly6k | NM_029627 | -4.982610477 |
| Slc13a5 | NM_001004148 | -2.661816794 |
| Rnf43 | NM_172448 | -1.270938463 |
| Cldn7 | NM_001193619 | -2.418945287 |
| Ms4a4d | NM_025658 | -1.26749296 |
| Cyp2c69 | NM_001104525 | -4.909594031 |
| Dixdc1 | NM_178118 | -1.435879209 |
| Tmtc2 | NM_177368 | -2.02681834 |
| Dlg3 | NM_001177779 | -0.651749882 |
| Cyp2c40 | NM_010004 | -3.54742528 |
| Mafb | NM_010658 | -1.130493421 |

| | | |
|---|---|---|
| Chic1 | NM_009767 | -2.885500936 |
| Pear1 | NM_001289601 | -0.936063251 |
| Cbx4 | NM_007625 | -1.106175103 |
| Rtn1 | NM_001286448 | -1.342024669 |
| Mylk | NM_139300 | -0.933867965 |
| Rasef | NM_001017427 | -3.037818479 |
| Patz1 | NM_019574 | -0.848698942 |
| Bmp4 | NM_007554 | -1.276903728 |
| Nup210 | NM_018815 | -0.795538516 |
| Slc15a2 | NM_021301 | -1.781197853 |
| Prr12 | NM_175022 | -0.68586034 |
| Cnksr3 | NM_172546 | -0.788482769 |
| Nxpe2 | NM_030069 | -2.275259543 |
| Ldoc1l | NM_177630 | -1.748722999 |
| Cyp3a16 | NM_007820 | -5.54437442 |
| Prlr | NM_001253781 | -1.916975144 |
| Plcd3 | NM_152813 | -1.725069434 |
| Spint1 | NM_016907 | -2.24902339 |
| Ccl19 | NM_011888_1 | -1.777382218 |
| Scnn1a | NM_011324 | -2.50512956 |
| Pde2a | NM_001243758 | -0.606890804 |
| Slc10a1 | NM_011387 | -2.268984724 |
| Kmt2c | NM_001081383 | -0.983812281 |
| Cyp2c29 | NM_007815 | -2.192212174 |
| Fcgbp | NM_001122603 | -2.365431364 |
| Stard8 | NM_199018 | -0.758732368 |
| Gem | NM_010276 | -1.50711356 |
| Eng | NM_001146348 | -0.676154653 |
| Hgf | NM_001289461 | -1.167468373 |
| Enah | NM_001083120 | -2.362987034 |
| Abcd4 | NM_008992 | -0.947391804 |
| Pitpnm3 | NM_001081641 | -2.810969121 |
| Magi1 | NM_010367 | -0.714417736 |
| Irf2bp1 | NM_178757 | -0.70137882 |
| Cux2 | NM_007804 | -2.368833979 |
| Amigo1 | NM_001287093 | -1.515469739 |
| Lpin1 | NM_015763 | -1.342023364 |
| Dkk3 | NM_015814 | -1.535928984 |
| Pcdhgc3 | NM_033581 | -0.922182191 |
| Bbs1 | NM_001033128 | -1.540705771 |
| Erich5 | NM_173421 | -2.792000467 |
| Evpl | NM_025276 | -1.454836773 |
| Smg6 | NM_001002764 | -0.605136533 |
| Kif12 | NM_010616 | -2.758004818 |
| Arid1a | NM_001080819 | -0.650997032 |
| Aff3 | NM_001290814 | -1.783015791 |
| Glul | NM_008131 | -1.923106306 |

| | | |
|---|---|---|
| St3gal6 | NM_018784 | -1.191917558 |
| Epha7 | NM_010141 | -1.905428885 |
| Sntg1 | NM_027671 | -1.890063636 |
| Esrp1 | NM_194055 | -3.405258858 |
| Thra | NM_178060 | -0.693122315 |
| Scn7a | NM_009135 | -2.02242753 |
| Acaca | NM_133360 | -0.943802722 |
| Pcif1 | NM_146129 | -0.622952745 |
| Slco2b1 | NM_175316 | -1.719674641 |
| Hip1r | NM_145070 | -0.691879012 |
| Tgfbr3 | NM_011578 | -1.270537069 |
| 5930403L1 | NR_045643 | -3.216866773 |
| Cyp2d22 | NM_019823 | -1.081090369 |
| Tns1 | NM_027884 | -0.763318456 |
| Bcl9l | NM_030256 | -1.049356757 |
| Pfkfb1 | NM_008824 | -1.256942101 |
| Tbc1d9b | NM_029745 | -0.605808675 |
| Usp11 | NM_145628 | -1.499896854 |
| Ep400 | NM_029337 | -0.604329543 |
| Sgk2 | NM_013731 | -1.266779996 |
| Hnrnpa3 | NM_198090 | -0.696927811 |
| Galnt4 | NM_015737 | -0.753490115 |
| Kazald1 | NM_178929 | -2.138051845 |
| Ski | NM_011385 | -0.658705062 |
| Itgae | NM_008399 | -1.657552119 |
| Osbpl5 | NM_024289 | -1.244068242 |
| Dab1 | NR_104385 | -3.146452167 |
| Cdkn1c | NM_001161624 | -1.32275145 |
| Sp4 | NM_001166385 | -1.191546626 |
| Ptpru | NM_001083119 | -1.671849485 |
| Ttll10 | NM_029264 | -2.639709107 |
| Gpm6a | NM_153581 | -1.821597272 |
| Col4a6 | NM_053185 | -2.778787463 |
| Gm15800 | NM_181421 | -0.943694382 |
| Gabbr2 | NM_001081141 | -3.002889259 |
| 9130008F2 | NM_027834 | -1.838572706 |
| Zcchc3 | NM_175126 | -1.529783545 |
| Ugt1a5 | NM_201643 | -3.452121811 |
| Art4 | NM_026639 | -1.306306011 |
| Nfib | NM_001113209 | -0.674725461 |
| Tmem204 | NM_001001183 | -0.88436869 |
| Zfp354c | NM_013922 | -1.796669861 |
| Nfat5 | NM_001286260 | -0.974486551 |
| Sobp | NM_175407 | -2.535167593 |
| Tmem108 | NM_178638 | -2.086808965 |
| Gaa | NM_008064 | -0.672538149 |
| Pdgfrb | NM_001146268 | -1.498304131 |

| | | |
|---|---|---|
| Etohd2 | NR_015349 | -1.782401615 |
| Lrp1 | NM_008512 | -1.111559008 |
| A230001M | NR_040391 | -3.229822867 |
| BC024139 | NM_001142968 | -3.248741382 |
| Sgpp2 | NM_001004173 | -2.226826158 |
| Ptprb | NM_029928 | -0.962891748 |
| Sned1 | NM_172463 | -2.501468986 |
| Lrrn3 | NM_001271709 | -1.785445596 |
| Mllt6 | NM_139311 | -0.632156906 |
| Serpina5 | NM_172953 | -3.308294474 |
| 1700011H1 | NM_025956 | -2.692090089 |
| Wipf2 | NM_197940 | -0.601407434 |
| Olfm1 | NM_001038613 | -1.312128723 |
| D630045M | NR_045293 | -2.041744436 |
| Shank2 | NM_001081370 | -1.02975289 |
| Zfp395 | NM_199029 | -1.526316881 |
| Cacna1a | NM_001252060 | -1.506162061 |
| Cmah | NM_001284519 | -1.781899762 |
| Ajap1 | NM_001099299 | -2.093478488 |
| Gltscr1 | NM_001081418 | -0.811477025 |
| Gdf2 | NM_019506 | -1.604122085 |
| Neurl4 | NM_001291119 | -0.750958446 |
| Dzip1 | NM_025943 | -1.297042934 |
| Setd1b | NM_001040398 | -1.014265396 |
| Zfpm2 | NM_011766 | -1.81767398 |
| Nlgn2 | NM_198862 | -1.384025536 |
| Cdk19 | NM_198164 | -0.615330355 |
| Ets1 | NM_011808 | -0.571332084 |
| Sel1l | NM_011344 | -0.802208505 |
| Pnpla3 | NM_054088 | -3.056132802 |
| Rassf9 | NM_146240 | -2.589810844 |
| Sh3yl1 | NM_013709 | -0.647285788 |
| Amotl1 | NM_001081395 | -1.027263027 |
| Kmt2e | NM_026984 | -0.739586547 |
