## Supplementary Table S5-Gene Lists- Atg7HMGB1DKO only downregulated for "The HMGB1-RAGE axis modulates the growth of autophagy-deficient hepatic tumors"

**Table S5: List of downregulated genes in tumors of *Atg7-/-/Hmgb1-/-* liver**

| Name | Trxid | logFC.A7HDKO_TumvsA7HDKO_NT |
|---|---|---|
| Gngt1 | NM_010314 | -2.991671437 |
| 0610005C13Rik | NR_038165 | -1.37674218 |
| 4833411C07Rik | NR_102285 | -1.444868642 |
| Oxld1 | NM_025560 | -1.652009523 |
| Coq10a | NM_001081040 | -1.283993202 |
| Fndc4 | NM_022424 | -1.324545036 |
| Magix | NR_037581 | -1.612042172 |
| 3930402G23Rik | NR_030715 | -2.802959557 |
| Rnf186 | NM_025786 | -1.719867592 |
| Macrod1 | NM_134147 | -1.48211712 |
| Mlycd | NM_019966 | -1.175649095 |
| Immp2l | NM_053122 | -1.39073284 |
| Paqr9 | NM_198414 | -1.742957219 |
| Igf1 | NM_001111276 | -1.497148573 |
| Folr2 | NM_001303239 | -1.310857436 |
| Acot13 | NM_025790 | -1.090688869 |
| Mtfp1 | NM_026443 | -1.519066821 |
| Pbld1 | NM_026701 | -1.153013545 |
| Urah | NM_029821 | -1.349849772 |
| Ccs | NM_016892 | -1.207359814 |
| Spc24 | NM_026282 | -1.69812334 |
| Pdzk1 | NM_001146001 | -1.098796779 |
| Nit2 | NM_023175 | -1.249207029 |
| Sucnr1 | NM_032400 | -1.128769832 |
| Id3 | NM_008321 | -1.212305302 |
| Hes6 | NM_019479 | -1.072919047 |
| Smlr1 | NM_001195596 | -1.137246806 |
| C8b | NM_133882 | -1.330941687 |
| Clec4g | NM_029465 | -1.269851222 |
| Apoc4 | NM_007385 | -1.867126759 |
| Mrpl41 | NM_001031808 | -1.232975888 |
| Tprkb | NM_176842 | -0.918740282 |
| Clec4f | NM_016751 | -1.351916473 |
| 1110001J03Rik | NM_025363 | -1.623143987 |
| Ppp1r3g | NM_029628 | -2.208744803 |
| Tmem256 | NM_026982 | -1.635748198 |
| Clec2d | NM_053109 | -1.259688098 |
| Ndufa5 | NM_026614 | -1.074021176 |
| Pyurf | NM_025574 | -1.007449576 |
| Enpep | NM_007934 | -1.039835448 |
| Ggact | NM_145466 | -1.173081823 |
| Glyat | NM_145935 | -0.9503296 |
| Ces1e | NM_133660 | -1.907378795 |
| Fuom | NM_001286218 | -1.237989955 |
| Bphl | NM_026512 | -0.827886585 |
| Tlcd1 | NM_001291236 | -1.14240794 |
| Lipc | NM_008280 | -1.539190053 |

| | | |
|---|---|---|
| Paox | NM_153783 | -1.156991255 |
| Wfdc1 | NM_023395 | -2.002798463 |
| Fh1 | NM_010209 | -1.126233806 |
| S100a1 | NM_011309 | -1.210282944 |
| A430005L14Rik | NM_175287 | -1.250686155 |
| Oaf | NM_178644 | -1.295871963 |
| Inmt | NM_009349 | -1.795965066 |
| Mustn1 | NM_181390 | -1.574355374 |
| Iah1 | NM_026347 | -1.174987304 |
| Rpain | NM_027186 | -1.323095971 |
| Ndufb2 | NM_026612 | -1.08658584 |
| Gstt2 | NM_010361 | -1.041854623 |
| 1810008I18Rik | NR_045301 | -0.873553785 |
| Bnip3 | NM_009760 | -0.96194259 |
| Uqcr11 | NM_025650 | -1.103506863 |
| Slc22a30 | NM_177002 | -1.051531892 |
| D130043K22Rik | NM_001081051 | -1.446848745 |
| Gstz1 | NM_001252556 | -1.294356127 |
| Fbp1 | NM_019395 | -1.40421119 |
| Ces1d | NM_053200 | -1.648764442 |
| Gm5617 | NM_001004191 | -1.458607724 |
| Dnajc15 | NM_025384 | -1.217761631 |
| Cdadc1 | NM_001168538 | -0.817471556 |
| Chchd10 | NM_175329 | -1.491896924 |
| Wnt5b | NM_009525 | -1.412488204 |
| Pdk1 | NM_172665 | -1.105232445 |
| Igfals | NM_008340 | -1.163630735 |
| Uqcrq | NM_025352 | -1.50573235 |
| Hebp1 | NM_013546 | -1.220599634 |
| Hagh | NM_024284 | -0.974056252 |
| Sirt3 | NM_022433 | -0.959511674 |
| Hpd | NM_008277 | -1.47774132 |
| Uros | NM_009479 | -0.885436045 |
| Ndufaf6 | NM_001085493 | -1.161418905 |
| Fggy | NM_001113412 | -1.149279823 |
| Plk3 | NM_013807 | -1.523218922 |
| Tmem14c | NM_025387 | -1.106940152 |
| Tmem261 | NM_025849 | -1.185568691 |
| Zfyve21 | NM_026752 | -1.080947326 |
| Agmat | NM_001081408 | -1.687638132 |
| Nudt6 | NM_153561 | -1.058693702 |
| Ndufa3 | NM_025348 | -0.97233289 |
| Pcyt2 | NM_024229 | -1.066340987 |
| Msrb1 | NM_013759 | -1.165880687 |
| Ndufb7 | NM_025843 | -1.202203986 |
| Qdpr | NM_024236 | -1.343795337 |
| Acbd4 | NM_025988 | -1.003643824 |
| AI317395 | NM_144821 | -1.507808559 |
| Inhbc | NM_010565 | -1.112767746 |

| | | |
|---|---|---|
| 1700001C19Rik | NM_029296 | -1.062068661 |
| Mmaa | NM_133823 | -0.869854932 |
| D130020L05Rik | NR_038048 | -1.273544209 |
| H2-Ke6 | NM_013543 | -1.066588619 |
| Ndufs7 | NM_029272 | -0.987836844 |
| Deb1 | NM_026794 | -1.096642395 |
| H2afv | NM_029938 | -1.007262754 |
| Gamt | NM_010255 | -1.134672685 |
| Uqcrh | NM_025641 | -1.182800574 |
| Adtrp | NM_001145875 | -1.092343224 |
| Atp5g1 | NM_007506 | -0.996770866 |
| Hamp | NM_032541 | -2.783920815 |
| Rab30 | NM_029494 | -0.992121501 |
| Tmem219 | NM_026827 | -1.151804313 |
| 1810058I24Rik | NR_015608 | -0.948755967 |
| Aurkaip1 | NM_025338 | -1.083393647 |
| Dbi | NM_007830 | -1.299265987 |
| Lage3 | NM_025410 | -1.049252998 |
| Grcc10 | NM_013535 | -0.949085935 |
| Adgrv1 | NM_054053 | -1.947398605 |
| Gm4737 | NM_001304528 | -1.287206631 |
| Pcbd2 | NM_028281 | -0.893246913 |
| Zfand2b | NM_001159905 | -1.060674902 |
| 1700012D01Rik | NR_045171 | -2.002798307 |
| Mrpl34 | NM_053162 | -1.052335056 |
| Stra13 | NM_016665 | -1.17343711 |
| 2810428I15Rik | NM_025577 | -1.049688039 |
| 2310010J17Rik | NR_046007 | -1.177314371 |
| Tmed1 | NM_010744 | -0.907149367 |
| Ddt | NM_010027 | -1.162212055 |
| Sar1b | NM_025535 | -1.016896361 |
| Khdrbs3 | NM_010158 | -1.187500225 |
| Fxn | NM_008044 | -1.006884199 |
| BC024386 | NR_015583 | -1.751708905 |
| Pex7 | NM_001161825 | -0.743869486 |
| 1110032A03Rik | NM_023483 | -0.814029941 |
| Suox | NM_173733 | -0.94755068 |
| Tefm | NM_183275 | -0.954912383 |
| Ccdc107 | NM_001037913 | -1.192865818 |
| Ccdc148 | NM_001001178 | -1.319331056 |
| Phyhd1 | NM_001281829 | -0.839888201 |
| Marc1 | NM_001290273 | -1.083780102 |
| Pxmp2 | NM_008993 | -1.150013775 |
| Cyp2d13 | NR_003552 | -2.328179377 |
| Apoa2 | NM_001305550 | -1.834361418 |
| Ict1 | NM_026729 | -1.022554824 |
| Fam25c | NM_183278 | -1.727379682 |
| Dhrs4 | NM_001037938 | -1.146408779 |
| Cecr5 | NM_144815 | -0.837142907 |

| | | |
|---|---|---|
| Mgmt | NM_008598 | -1.015176713 |
| 2310039H08Rik | NM_025966 | -1.26826527 |
| Ndufs8 | NM_001271443 | -1.011725497 |
| Atp6v0b | NM_033617 | -1.047797866 |
| Dgcr6 | NM_010047 | -1.400963563 |
| Slc2a4 | NM_009204 | -2.145090731 |
| Ecsit | NM_001253897 | -0.877414691 |
| Uqcr10 | NM_197979 | -1.108866267 |
| Pet100 | NM_001195244 | -1.119783973 |
| Ech1 | NM_016772 | -1.074591456 |
| Gcdh | NM_001044744 | -1.093291241 |
| Chchd1 | NM_025366 | -1.057415377 |
| Smim4 | NM_001308464 | -1.488565389 |
| Ndufb6 | NM_001033305 | -0.942043581 |
| Tmem205 | NM_001253867 | -1.024937522 |
| Bola1 | NM_026975 | -1.030460922 |
| Mif4gd | NM_001243587 | -0.904209303 |
| Morn2 | NM_194269 | -1.064444849 |
| Uqcrb | NM_026219 | -0.930849023 |
| Il11ra1 | NM_001163401 | -1.039542879 |
| 1700048O20Rik | NR_033553 | -2.184090035 |
| Csad | NM_144942 | -0.981699632 |
| Pigyl | NM_001082532 | -0.886736668 |
| Mvb12a | NM_028617 | -0.814578425 |
| Tmem223 | NM_025791 | -1.078931389 |
| Gcsh | NM_026572 | -0.85453671 |
| Tmem53 | NM_029589 | -1.03177916 |
| Cib2 | NM_019686 | -1.298930386 |
| Tmem11 | NM_173453 | -0.871986574 |
| Znhit2 | NM_013859 | -0.888668155 |
| 4930581F22Rik | NR_029475 | -1.224454876 |
| Il1rap | NM_001159317 | -1.145009994 |
| Uox | NM_009474 | -1.639771737 |
| Ndufb8 | NM_026061 | -0.871193029 |
| Rbks | NM_153196 | -0.900749538 |
| Ndufs4 | NM_010887 | -0.809790039 |
| Fbxl15 | NM_133694 | -1.179188003 |
| Mtrf1 | NM_145960 | -0.935067764 |
| Ndufb11 | NM_019435 | -0.873197087 |
| D10Jhu81e | NM_138601 | -0.83969397 |
| Phpt1 | NM_029293 | -0.918335455 |
| Fxyd1 | NM_019503 | -1.226103632 |
| 2010107E04Rik | NM_027360 | -1.051488558 |
| Timmdc1 | NM_024273 | -0.907044558 |
| Gcat | NM_001161712 | -0.968599817 |
| Dcxr | NM_026428 | -1.222706874 |
| Cd59a | NM_007652 | -1.062211685 |
| Leap2 | NM_153069 | -2.348168131 |
| Etfb | NM_026695 | -1.070383539 |

| | | |
|---|---|---|
| 2010012O05Rik | NM_025563 | -0.853132495 |
| Ndufc2 | NM_024220 | -0.785693826 |
| Echdc2 | NM_001254754 | -0.891794931 |
| 1700096K18Rik | NR_027388 | -1.370827496 |
| Gstk1 | NM_029555 | -0.987831357 |
| Adra1b | NM_001284381 | -1.152113945 |
| Rsrp1 | NM_023665 | -0.816494644 |
| Timm13 | NM_013895 | -0.799784014 |
| Slc23a1 | NM_011397 | -0.84579558 |
| Nudt7 | NM_001290180 | -2.027964575 |
| Gja4 | NM_008120 | -0.962595741 |
| Cradd | NM_009950 | -0.870986449 |
| Pemt | NM_001290011 | -1.979108405 |
| Fam173a | NM_001285984 | -0.830410435 |
| Timm9 | NM_001286203 | -0.826735888 |
| Dpm3 | NM_026767 | -1.113568663 |
| Ifi27l2a | NM_001281830 | -1.253732955 |
| Acot4 | NM_134247 | -1.603216302 |
| Lcn8 | NM_033145 | -2.934856154 |
| St3gal4 | NM_009178 | -0.744423407 |
| Gadd45gip1 | NM_183358 | -0.8966717 |
| Gm10658 | NR_045886 | -1.090281568 |
| Ptgr2 | NM_029880 | -0.767205576 |
| Cml1 | NM_023160 | -1.978211435 |
| Lin7a | NM_001284329 | -1.521081991 |
| Nr1h4 | NM_001163504 | -0.906153418 |
| Ndufb3 | NM_025597 | -0.885636622 |
| LOC100504703 | NR_040660 | -1.332841579 |
| Ndufa1 | NM_019443 | -0.889942346 |
| Mrpl12 | NM_027204 | -0.930926321 |
| Cox16 | NM_025461 | -1.389247142 |
| Rilp | NM_001029938 | -1.068485856 |
| Saysd1 | NM_026209 | -0.891265907 |
| Hddc3 | NM_026812 | -1.256116731 |
| Sult2a7 | NM_001184981 | -5.667202384 |
| Cdc26 | NM_139291 | -0.919963037 |
| Mipep | NR_040642 | -0.676231864 |
| Commd1 | NM_144514 | -1.076211244 |
| Pah | NM_008777 | -0.886441174 |
| Ebp | NM_007898 | -0.992554612 |
| Mrps33 | NM_010270 | -0.858744645 |
| Fpgs | NM_010236 | -0.920520341 |
| Ppp1r14a | NM_026731 | -1.233260994 |
| Mrpl32 | NM_029271 | -0.69943709 |
| 0610011F06Rik | NM_026686 | -1.243491019 |
| Coasy | NM_001305982 | -0.9102581 |
| Sdhb | NM_023374 | -0.917634129 |
| Mrpl27 | NM_053161 | -0.795763261 |
| Ndufa11 | NM_027244 | -0.869033723 |

| | | |
|---|---|---|
| Gm4952 | NM_001013762 | -0.960028088 |
| Aadac | NM_023383 | -0.927148556 |
| Coq9 | NM_026452 | -0.823819058 |
| Atoh8 | NM_153778 | -0.922166527 |
| Igfbp2 | NM_008342 | -1.402950668 |
| Isca2 | NM_028863 | -1.058289826 |
| Timm17b | NM_011591 | -0.878712122 |
| Pex11g | NM_026951 | -1.238882235 |
| 1110065P20Rik | NM_001142727 | -0.940468244 |
| Apoa1bp | NM_144897 | -0.898178813 |
| Phyh | NM_010726 | -1.290388157 |
| Cnpy2 | NM_019953 | -0.849132235 |
| Fcna | NM_007995 | -1.44075524 |
| Kxd1 | NM_029366 | -0.749356026 |
| Ndufa13 | NM_023312 | -0.793159751 |
| Ndufa6 | NM_025987 | -0.811040366 |
| Mrpl48 | NR_003559 | -0.770809012 |
| Atraid | NM_027855 | -0.859280741 |
| Arg1 | NM_007482 | -1.169788997 |
| Eif4a2 | NM_013506 | -0.788355625 |
| Pts | NM_011220 | -0.77370767 |
| Romo1 | NM_025946 | -0.933988579 |
| Cyp4f15 | NM_134127 | -1.270713854 |
| Il18 | NM_008360 | -1.038252575 |
| Slc25a10 | NM_013770 | -1.11137211 |
| Aass | NM_013930 | -0.799940791 |
| Agpat6 | NM_018743 | -0.843726942 |
| S100a13 | NM_009113 | -0.961599946 |
| Ift20 | NM_018854 | -1.144536691 |
| Grtp1 | NM_025768 | -0.733506846 |
| 2010003K11Rik | NM_027237 | -1.231059958 |
| Cox7a2 | NM_009945 | -0.828919859 |
| Fahd2a | NM_029629 | -0.877385032 |
| Ablim3 | NM_001164491 | -0.974090016 |
| Bst2 | NM_198095 | -0.834040277 |
| Naip1 | NM_008670 | -1.400984595 |
| Cmc2 | NM_026844 | -0.989389126 |
| Decr1 | NM_026172 | -1.212149395 |
| Aspdh | NM_026690 | -1.529996849 |
| Atp5o | NM_138597 | -1.006232494 |
| Higd2a | NM_025933 | -0.848734603 |
| Atp6v1g1 | NM_024173 | -0.757526868 |
| Aamdc | NM_183251 | -0.71251325 |
| Minos1 | NM_001163006 | -0.894912409 |
| Dhrs1 | NM_026819 | -1.04842775 |
