## Supplementary Table S7-List of Primers used for qPCR for "The HMGB1-RAGE axis modulates the growth of autophagy-deficient hepatic tumors"

**Supporting Table 7.List of Primers used for qPCR**

| Gene name | Forward primer | Reverse Primer |
|---|---|---|
| Actin | 5'-ACTATTGGCAACGAGCGGTT-3' | 5'-CAGGATTCCATACCCAAGAAGGA-3' |
| Angpt1 | 5'-CTGTGGCCCTTCCAATCTAAA-3' | 5'-GTAACTGGGCCCTTTGAAGTAG-3' |
| Angpt2 | 5'-ACAGCTGTGATGATAGAGATTGG-3' | 5'-CGAGTCTTGTCGTCTGGTTTAG-3' |
| Cd133 | 5'-GAAAAGTTGCTCTGCGAACC-3' | 5'-TCTCAAGCTGAAAAGCAGCA-3' |
| Cd200 | 5'-GGGCAGTCTGGTATTCAGGA-3' | 5'-CTGGGTCACCACTTCCACTT-3' |
| Cd24a | 5'-CTTCTGGCACTGCTCCTACC-3' | 5'-GAGAGAGAGCCAGGAGACCA-3' |
| Cd34 | 5'-GGGTAGCTCTCTGCCTGATG-3' | 5'-CAGTTGGGGAAGTCTGTGGT-3' |
| Cd4 | 5'-AGGAAGTGAACCTGGTGGTG-3' | 5'-CTCCTGCTTCAGGGTCAGTC-3' |
| Cd44 | 5'-TGGATCCGAATTAGCTGGAC-3' | 5'-AGCTTTTTCTTCTGCCCACA -3' |
| Cd8 | 5'-TATGGCTTCATCCCACAACA-3' | 5'-GACTGGCACGACAGAACTGA-3' |
| Cd90 | 5'-CGCTCTCCTGCTCTCAGTCT-3' | 5'-GTTATTCTCATGGCGGCAGT-3' |
| Cyclin A | 5'-ACAGAGCTGGCCTGAGTCAT-3' | 5'-TTGACTGTTGGGCATGTTGT-3' |
| Cyclin B | 5'-AGCGAAGAGCTACAGGCAAG-3' | 5'-TCACACACAGGCACCTTCTC-3' |
| Cyclin D | 5'-GCGTACCCTGACACCAATCT-3' | 5'-ATCTCCTTCTGCACGCACTT-3' |
| F4/80 | 5’-TGCATCTAGCAATGGACAGC-3’ | 5’-GCCTTCTGGATCCATTTGAA-3’ |
| Gstm1 | 5′-CTACCTTGCCCGAAAGCAC-3′ | 5′-ATGTCTGCACGGATCCTCTC-3′ |
| Hmgb1 | 5'-CGCGGAGGAAAATCAACTAA-3' | 5'-GCAGACATGGTCTTCCACCT-3' |
| Il-17 | 5’-CTGGAGGATAACACTGTGAGAGT-3’ | 5’-TGCTGAATGGCGACGGAGTTC-3’ |
| Il-1b | 5’-GCTGCTTCCAAACCTTTGAC -3’ | 5’-TGTCCTCATCCTGGAAGGTC -3’ |
| Il-6 | 5'-AGTTGCCTTCTTGGGACTGA-3' | 5'-TCCACGATTTCCCAGAGAAC-3' |
| Klf4 | 5'-CTGAACAGCAGGGACTGTCA-3' | 5'-GAGGGGACTTGTGACTGCAT-3' |
| Ly6a | 5'-CTCAGGAGGCAGCAGTTATT-3' | 5'-GTACCCAGGATCTCCATACTTTC-3' |
| Ly6c1/Gr1 | 5'-AGTCCTGTGTGCTCATTCTTC-3' | 5'-AGTCTCAATTGGCACTCCATAG-3' |
| Ly6d | 5'-AAACCGTCACCTCAGTGGAG-3' | 5'-CATAGGTCAGTCTGGCAGCA-3' |
| Nanog | 5'-ACCCAACTTGGAACAACCAG-3' | 5'-GAAGTTATGGAGCGGAGCAG-3' |
| Nqo1 | 5′-AGCGTTCGGTATTACGATCC-3′ | 5′-AGTACAATCAGGGCTCTTCTCG-3′ |
| Oct4 | 5'-CACGAGTGGAAAGCAACTCA-3' | 5'-TTCATGTCCTGGGACTCCTC-3' |
| Pcna | 5'-ATCCTGAAGAAGGTGCTGGA -3' | 5'-TGAGACGAGTCCATGCTCTG-3' |
| Pdgfb | 5'-GACTACCTGCACCGGAACAA-3' | 5'-GTGCAACATGGGCACGTAA-3' |
| Sox2 | 5'-AACGCCTTCATGGTATGGTC-3' | 5'-CGGACAAAAGTTTCCACTCC-3' |
| Tnfa | 5′-CGTCAGCCGATTTGCTATCT-3′ | 5′-CGGACTCCGCAAAGTCTAAG-3′ |
| Vegfa | 5'-CAGGCTGCTGTAACGATGAA-3' | 5'-TTTCTTGCGCTTTCGTTTTT-3' |
