## Supplementary Table S8-List of Antibodies used for immunostaining and western blot for "The HMGB1-RAGE axis modulates the growth of autophagy-deficient hepatic tumors"

**Supporting Table 8. List of Antibodies used for immunostaining and western blot**

| Antibody/Species | Source/Catalogue Number/Dilution |
|---|---|
| ACTIN/Mouse | Sigma/5441/1:5000 |
| Alexa Fluor 488 goat anti-rabbit | Molecular Probes/A11034/1:250 |
| ATG7/Rabbit | Cell Signal/#2631/1:1000 |
| CD3/Rat | BD Pharmagen/#557306/1:100 |
| CD45R/Rat | BD Pharmagen/#553089/1:100 |
| CK19/Rat | Developmental Studies Hybridoma Bank/#1DB-001-0000868971/1:200 |
| Cy3-Anti Rat | Jackson ImmunoResearch Laboratories Inc/#712-165-150/1:500 |
| Cy3-Anti-Goat | Jackson ImmunoResearch Laboratories Inc/#705-165-147 /1:250 |
| Cy3-Anti-Mouse | Jackson ImmunoResearch Laboratories Inc/#115 165 146 /1:250 |
| CYCLIN D/Mouse | Santa Cruz/#sc-450/1:100 |
| CYCLIN E/Rabbit | Santa Cruz/#sc-481/1:100 |
| DESMIN/Rabbit | Thermo/#RB9014P0/1:200 |
| F4/80/Rat | Bio-Rad/# MCA497G/1:100 |
| GAPDH/Mouse | Novus/#NB300-21/1:3000 |
| GSK-3β/Rabbit | Cell Signal/#12456/1:1000 |
| HMGB1/Rabbit | Abcam/#ab18256/1:1000 |
| HNF4a/Goat | Santa Cruz/#sc-6556/1:100 |
| HRP-labeled Mouse secondary antibody | Jackson ImmunoResearch Laboratories Inc/#115-035-062/1:5000 |
| HRP-labeled Rabbit secondary antibody | Jackson ImmunoResearch Laboratories Inc/#111-035-045/1:5000 |
| LC3B/Rabbit | Sigma/#L7543/1:1000 |
| NQO1/Rabbit | Abcam/#ab34173/1:3000 |
| NRF2/Rabbit | Cell Signal/#12721/1:1000 |
| PCNA/Mouse | Cell Signal/#2586/1:1000 |
| Phospho-AKT (Thr308)/Rabbit | Cell Signal/#13038/1:1000 |
| Phospho-GSK-3β(Ser9)/Rabbit | Cell Signal/#5558/1:1000 |
| Phospho-PDK1 (Ser24)/Rabbit | Cell Signal/#3438/1:1000 |
| Phospho-SAPK/JNK (Thr183/Tyr185) | Cell Signal/#9251/1:1000 |
| SOX9/Rabbit | EMB Millipore/#AB5535/1:1000 |
| SQSTM1/Mouse | Abnova/#H00008878-M01/1:1000 |
| Total-AKT/Rabbit | Cell Signal/#4691/1:1000 |
| Total-PDK1/Rabbit | Cell Signal/#3062/1:1000 |
| Total-SAPK/JNK | Cell Signal/#9252/1:1000 |
| UBIQUITIN/Rabbit | Santa Cruz/#sc-9133/1:100 |
| Phospho-c-JUN (Ser63)/Rabbit | Cell Signal/#9261/1:1000 |
| Total-JUN/Rabbit | Santa Cruz/#sc-30055/1:100 |
| Ki67/Rabbit | Abcam/#ab15580/1:3000 |
| Phospho-4E-BP1 (Thr37-46)/Rabbit | Cell Signal/#9459/1:1000 |

| | |
|---|---|
| Total 4E-BP1/Rabbit | Cell Signal/#9452/1:1000 |
| Phospho-p70 S6 kinase (Thr389)/Mouse | Cell Signal/#9234/1:1000 |
| Total S6 Kinase/Rabbit | Cell Signal/#9202/1:1000 |
| Phospho-S6/Rabbit | Cell Signal/#2211/1:1000 |
| Total S6 /Rabbit | Cell Signal/#2217/1:1000 |
| Total mTOR/Rabbit | Cell Signal/#2983/1:1000 |
| Phospho AMPKα (Thr172)/Rabbit | Cell Signal/#2531/1:1000 |
| Total-AMPKα (Thr172)/Rabbit | Cell Signal/#2532/1:1000 |
| Phospho-ERK/MAPK/Rabbit | Cell Signal/#4370/1:1000 |
| Total-ERK/MAPK/Rabbit | Cell Signal/#9102/1:1000 |
| Phospho-STAT3/Mouse | Santa Cruz/#sc-8059/1:100 |
| Total-STAT3/Mouse | Santa Cruz/#sc-8019/1:100 |
